# Regulatory T-cells in multiple sclerosis produce IL-10 in the central 1 nervous system but are activated by Epstein-Barr Virus

**DOI:** 10.1101/2024.07.30.605745

**Authors:** N. Pulvirenti, C. Righetti, F. Clemente, B. Serafini, C. Cordiglieri, C. Vasco, S. Maioli, A. Pietroboni, E. Galeota, B. Rosicarelli, C. Iannone, M. de Riz, A. Espadas de Arias, T. De Feo, L. Valenti, D. Prati, S. Abrignani, M. Gerosa, R. Caporali, D. Galimberti, E. Scarpini, J. Geginat

**Affiliations:** Fondazione Istituto Nazionale di Genetica Molecolare, Milan, Italy; Department of Neuroscience, Istituto Superiore di Sanità, Rome, Italy; Neurology-Neurodegenerative Diseases Unit, Fondazione IRCCS Ca’ Granda Ospedale Maggiore Policlinico, Milan, Italy; Laboratorio di Immunologia dei Trapianti - S.C. Trapianti Lombardia-NITp - Fond. IRCCS Cà Granda Ospedale Maggiore Policlinico, Milan, Italy; Department of Pathophysiology and Transplantation, Università degli Studi di Milano, Italy; Transfusion Medicine, Fondazione IRCCS Ca’ Granda Ospedale Maggiore Policlinico, Milano, Italy; Department of Clinical Sciences and Community Health, Università degli Studi di Milano, Italy; Centro Specialistico Ortopedico Traumatologico Gaetano Pini - CTO, Milan, Italy; Department of Biomedical, Surgical and Dental Sciences, Università degli Studi di Milano, Italy

## Abstract

Regulatory T-cells (Tregs) maintain immune homeostasis, but antigens activating adaptive Tregs in human pathologies are ill-defined. EOMES^+^type-1 regulatory (Tr1)-like T-cells had a dysregulated homeostasis in multiple sclerosis (MS), which was related to their activation in the central nervous system (CNS). EOMES^+^Tr1-like cells were enriched and clonally expanded in patient’s cerebrospinal fluid (CSF) and were the major IL-10-producing T-cell subset in the CNS. Regulatory T-cells from PwMS produced IL-10 and IFN-γ with antigens derived from Epstein-Barr Virus (EBV), but not from myelin. EOMES^+^Tr1-like cells responded selectively to the latency-associated antigen EBNA1, whereas FOXP3^+^Tregs and Th1-cells responded also to lytic EBV antigens. EBNA1-specific EOMES^+^Tr1-like cells were present in patients carrying the HLA-DRB1*15:01 risk allele, were associated with anti-EBNA1 IgG and disappeared upon therapeutic B-cell depletion. IL-10^+^EOMES^+^Tr1-like were present in MS brain lesions, and some were in the vicinity of EBV-infected B-cells and CD8^+^T-cells. Notably, EOMES^+^Tr1-like cells suppressed CD8+T-cell activation by EBV-infected B-cells. We propose that the insufficient protective functions of Tregs in MS are due to their aberrant anti-viral specificities that promote immune escape of a disease-associated virus.

## Introduction

Natural regulatory T-cells (Tregs) are generated in the thymus, express the transcription factor FOXP3 and are required to prevent autoimmunity (Ochs et al., 2007; Sakaguchi et al., 2006). A reduced number or functionality of FOXP3^+^Tregs to control the activation of autoreactive T-cells is therefore thought to play a crucial role in autoimmune diseases (Buckner, 2010). FOXP3^+^Tregs can however also be generated in the periphery in response to foreign antigens, which may be derived from pathogens (Suffia et al., 2006). Some regulatory T-cells lack FOXP3 expression but produce high levels of the anti-inflammatory cytokine IL-10 (Roncarolo et al., 2006). These IL-10 producing “type 1 regulatory T (Tr1)-cells” are adaptive Tregs, which are generated in the periphery in response to self- or foreign antigens (Haringer et al., 2009; Parish et al., 2014), and require thus TCR stimulation by specific antigens for their functions (Gagliani et al., 2010). Tr1-cells are highly heterogeneous (Brockmann et al., 2018). They may produce IL-10 alone or together with IFN-γ (Anderson et al., 2007; Jankovic et al., 2007; Jeon et al., 2012; O’Garra and Vieira, 2007) and inhibit excessive immune responses and immunopathology (O’Garra and Vieira, 2007). During persistent viral infections, IL-10 and IFN-γ co-producing Tr1-like cells are generated upon chronic antigenic stimulation and inhibit collateral tissue damage, but also viral clearance (Murata et al., 2021; Parish et al., 2014). In an increasing number of pathologies, including chronic viral infections, IL-10 and IFN-γ co-producing Tr1-like cells were reported to express the transcription factor EOMESODERMIN (EOMES) and the cytotoxic molecule Granzyme-K (GzmK) (Bonnal et al., 2021; Geginat et al., 2023; Gruarin et al., 2019; Haringer et al., 2009; Thelen et al., 2023; Zhang et al., 2017). Similar to FOXP3^+^Tregs, EOMES^+^Tr1-like cells can suppress CD8^+^T-cell responses (Bonnal et al., 2021; Pulvirenti et al., 2024), the key effector cells that control viral infections. Not all EOMES-expressing CD4^+^T-cells are however Tr1-like cells, since those expressing CCR6 or GzmB have pro-inflammatory properties (Dejean et al., 2019; Mazzoni et al., 2018; Pulvirenti et al., 2024). However, pro-inflammatory EOMES^+^Th1-cells may differentiate to anti-inflammatory EOMES^+^Tr1-like cells upon chronic antigenic activation in a negative feedback loop to inhibit immunopathology (Gabrysova et al., 2009; Geginat et al., 2023; O’Garra and Vieira, 2007)(Zhang, Immunity, 2025).

Multiple sclerosis (MS) is the most common chronic inflammatory demyelinating disease of the central nervous system (CNS) with a prevalent relapsing-remitting (RR-MS) onset that can evolve into a progressive course (P-MS). The aetiology is still unclear, but both genetic and environmental factors contribute critically to the overall disease risk (Olsson et al., 2017). The strongest genetic risk factor is the human leukocyte antigen (HLA) class II allele HLA-DRB1*15:01 which encodes an MHC class 2 molecule that could present critical self-peptides to activate autoreactive CD4^+^T-cells (Wang et al., 2020). Among environmental factors, infection with the almost ubiquitous herpesvirus Epstein-Barr Virus (EBV) is a necessary, although not sufficient, condition for initiating inflammatory brain damage leading to MS (Bjornevik et al., 2023). EBV-associated infectious mononucleosis and anti-Epstein Barr Virus nuclear antigen (EBNA)1 IgG levels interact with HLA-DRB1*15:01 to increase the risk of MS. However, the mechanism by which EBV promotes MS is still a focus of intense research (Aloisi et al., 2023; Christensen, 2006; Geginat et al., 2017; Murata et al., 2021). Accumulating evidence indicates an altered immune response to EBV in MS, and anti-EBV immune surveillance may be inefficient in MS (Geginat et al., 2017; Kuchroo and Weiner, 2022).

Regulatory T-cells are thought to be reduced or defective in MS, since they fail to prevent CNS damage. MS-like disease symptoms were however not reported in FOXP3-deficient mice or patients (Ochs et al., 2007), and reduced FOXP3^+^Treg numbers are not consistently associated with MS (Noori-Zadeh et al., 2016). FOXP3^+^Tregs proliferate however less upon TCR stimulation in MS (Carbone et al., 2014), produced higher amounts of IFN-γ and lower amounts of IL-10, and may thus be functionally impaired (Dominguez-Villar et al., 2011; Sumida et al., 2018). Tr1-like cells producing IL-10 and IFN-γ were shown to play a protective role in experimental autoimmune encephalomyelitis (EAE), a widely used animal model of autoimmunity-driven CNS inflammation (Heinemann et al., 2014). Moreover, the generation of IL-10 producing CD4^+^T-cells *in vitro* was defective in MS (Astier et al., 2006), suggesting that the differentiation of potentially protective Tr1-cells may be impaired. However, if EOMES^+^Tr1-like cells are involved in MS or anti-EBV immune control is unknown (Geginat et al., 2023). Notably, EOMES expression in CD4^+^T-cells promoted CNS inflammation in EAE (Stienne et al., 2016), suggesting that pro-inflammatory EOMES^+^T-cells could promote CNS damage. Moreover, cytotoxic CD4^+^EOMES^+^GzmB^+^T lymphocytes (CTL) were proposed to play a pathogenic role in MS (Peeters et al., 2017; Raveney et al., 2021). Here, we investigated the features of CD4^+^EOMES^+^T-cell subsets in MS. We found that EOMES^+^Tr1-like cells were activated *in vivo,* produced high amounts of IL-10, accumulated and were clonally expanded in the CNS. However, they were specific for the EBV-derived antigen EBNA1, suggesting that they are insufficient to prevent CNS autoimmune damage and promote EBV immune escape.

## Results

### The homeostasis of EOMES^+^Tr1-like cells is dysregulated in MS

T-cells from peripheral blood (PB) of recently diagnosed, treatment-naive RR-PwMS and age and sex-matched healthy donors (HDs, Table 1) were monitored by multidimensional flow cytometry. Bioinformatic analysis based on unsupervised clustering and dimensionality reduction algorithm, identified 27 T-cell clusters, including 12 CD4^+^ clusters (Figure 1A, sFigure 1). Cluster 11 displayed the phenotype of EOMES^+^Tr1-like cells (i.e. CD4^+^EOMES^+^GzmK^+^ CCR5^+^PD1^+^CD27^+^CCR6^−^ (Alfen et al., 2018; Geginat et al., 2023; Gruarin et al., 2019)), and co-clustered with CD4^+^EOMES^+^T-cells that expressed GzmK and CCR6 (Cluster (Cl) 10, sFigure 1C). In the Uniform Manifold Approximation and Projection (UMAP) maps (Figure 1A) these two GzmK^+^CD4^+^T-cell clusters were positioned in the centre of Th1-like clusters (Cl 5,6,8) and CD4^+^GZMB^+^CTL (Cl 12), consistent with an intermediate differentiation stage (De Simone et al., 2021; Pulvirenti et al., 2024). Surprisingly, we detected however no major changes in the composition of the T-cell compartments of PwMS and HDs (sFigure 1D).

**Figure 1:**
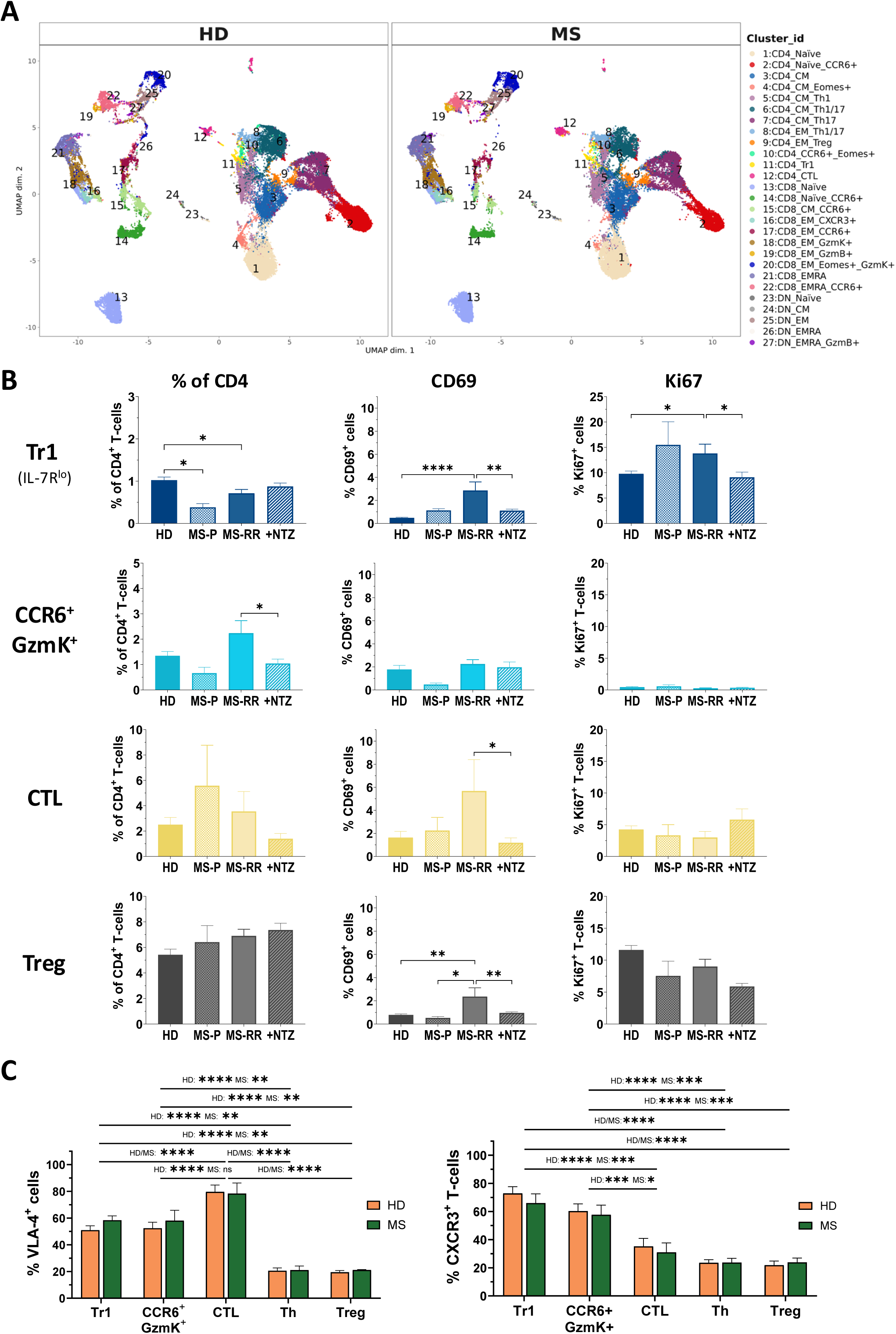
Dysregulated Homeostasis of EOMES^+^Tr1-like effector cells in PwMS. **A.** UMAP of T-cell clusters identified in peripheral blood of treatment-naïve RR-PwMS (n=14) and healthy donors (n=12) by multiparametric flow cytometry. Clusters are identified by numbers, different colors and named as indicated in the panel. **B.** Frequencies among CD4^+^T-cells, CD69 and Ki67 expression of the indicated T-cell subsets in HDs (n=35) untreated P-PwMS (n=3), treatment-naïve (n=11) and Natalizumab-treated RR-PwMS (n=20). **C.** VLA-4 and CXCR3 expression on CD4^+^T-cell subsets in treatment-naive RR-MS patients (n=4) and HDs (n=9). **B/C** Statistical significance was calculated by One-way ANOVA.

**Table 1:** Characteristics of PwMS and healthy donors.

|  | Healthy Donors | Treatment naive RR-MS | NTZ-treated RR-MS | Untreated P-MS | OCR-treated MS | Untreated MS (OCR Ctr) |
| --- | --- | --- | --- | --- | --- | --- |
| **Number of subjects** | 78 | 13 | 22 | 4<br>(3PP, 1SP) | 17 (8 PP, 3 SP, 6 RR) | 13<br>(3 PP, 10 RR) |
| **Female/male (ratio)** | 47/31<br>(1.5) | 7/6<br>(1.2) | 14/10<br>(1.4) | 0/4<br>(0) | 7/10<br>(0.7) | 5/8<br>(0.6) |
| **Age, median (range)** | 34<br>(20-61) | 34<br>(20-51) | 35<br>(20-62) | 48<br>(28-69) | 47<br>(25-69) | 44<br>(20-69) |
| **EDSS T0, median (range)** | n.a. | 1.0<br>(0.0-2.0) | 2.0<br>(1.0-5.5) | 2.8<br>(2.5-4.5) | 3.5<br>(0.0-6.5) | 1.0<br>(0.0-4.5) |
| **EDSS T1 (2023), median (range)** | n.a. | 1.0<br>(0.0-2.5) | 2.0<br>(1.0-6.0) | 3.0<br>(2.5-4.5) | 3.5<br>(0.0-7.0) | 1.5<br>(0.0-4.5) |

EDSS: Expanded Disability Status Scale; MS: Multiple Sclerosis; NTZ: Natalizumab; OCR: Ocrelizumab; P: Progressive; PP: Primary Progressive; RR: Relapsing-Remitting; SP: Secondary Progressive
In addition, we collected blood from 17 SLE patients with and without CNS involvement.

We therefore analysed EOMES^+^CD4^+^T-cell subsets more precisely by manual gating in a higher number of subjects, exploiting an established gating strategy (Cossarizza et al., 2021; Gruarin et al., 2019). Thus, we distinguished CD4^+^EOMES^+^ subsets according to the expression of GzmB (“CD4^+^CTL”) or GzmK from FOXP3^+^Tregs and helper T-cells (“Th”), which lack FOXP3 and Granzyme expression (sFigure 2A, B). CD4^+^GzmK^+^T-cells were further subdivided into pro-inflammatory GzmK^+^CCR6^+^ cells and two Tr1-containing GzmK^+^CCR6^−^ populations that expressed different levels of IL-7R. These subsets displayed characteristic cytokine profiles in PwMS (sFigure 2C), as we previously reported in HDs (Pulvirenti et al., 2024). Notably, IL-7R^lo^GzmK^+^T-cells had a mature Tr1 profile, since they possessed the highest IL-10 producing capacities but had a low capacity to up-regulate CD40L (sFigure 2C). We then analysed the frequencies and the *in vivo* activation state of T-cell subsets according to the expression of the early activation marker CD69 and the proliferation marker Ki67 *ex vivo* (Figure 1B, sFigure 3A). IL-7R^lo^Tr1-like cells were significantly reduced in PwMS and expressed higher levels of CD69 and Ki67 in RR-PwMS, indicating an increased *in vivo* turnover (Figure 1B). Also IL-7R^+^Tr1-like cells (sFigure 3A), CD4^+^CTL and FOXP3^+^Tregs expressed increased levels of CD69 in RR-MS, but their frequencies and *in vivo* turnover rates were not significantly altered. Pro-inflammatory CCR6^+^GzmK^+^ cells were in contrast increased in RR-MS but maintained low CD69 and Ki67 expression (Figure 1B). Finally, Th-cells represented consistently the large majority of CD4^+^T-cells and remained largely CD69^−^ and quiescent in PwMS (sFigure 3A). Thus, only IL-7R^lo^Tr1-like cells had an increased proliferation in PwMS and were significantly decreased. However, in Natalizumab-treated PwMS frequencies and *in vivo* turnover rates of IL-7R^lo^Tr1-like cells were similar to those from HDs. Moreover, CD69 expression on all EOMES^+^Tr1-cells, as well as on FOXP3^+^Tregs and CD4^+^CTL, was significantly reduced in Natalizumab-treated as compared to untreated RR-PwMS (Figure 1B, sFigure 3A). Since Natalizumab blocks the adhesion receptor VLA-4 and consequently lymphocyte homing to the CNS, these finding suggested that the altered Tr1 homeostasis in MS is related to CNS homing (Kivisakk et al., 2009). Consistently, the majority of EOMES^+^Tr1-like cells, as well as of CCR6^+^GzmK^+^ cells and of CD4^+^CTL, expressed VLA-4 in HDs and in PwMS (Figure 1C), suggesting that all cytotoxic CD4^+^T-cell subsets possess a constitutive high CNS homing potential and can be efficiently targeted by Natalizumab. Conversely, only minor fractions of FOXP3^+^Tregs and Th-cells were VLA-4^+^. Moreover, most EOMES^+^Tr1-cells and CCR6^+^GzmK^+^ cells expressed also CXCR3 (Figure 1C), a chemokine receptor that is involved in CNS homing in PwMS (Geginat et al., 2017; Kendirli et al., 2023). Notably, CD4^+^CTL, FOXP3^+^Tregs and Th-cells expressed significantly lower levels of CXCR3. Furthermore, also most recently divided Ki67^+^Tr1-like cells were CXCR3^+^, whereas CXCR3 expression on GZMK^−^Ki67^+^T-cell subsets was significantly lower (sFigure 3D).

Overall, these findings indicate that the homeostasis of EOMES^+^Tr1-like cells is dysregulated in MS and suggest that this alteration may be related to CNS homing.

### Clonally expanded EOMES^+^Tr1-like accumulate in the cerebrospinal fluid of PwMS

In order to investigate if EOMES^+^Tr1-like cells could be recruited to the CNS in PwMS, we interrogated a single cell RNA and TCR sequencing dataset of antigen-experienced CD4^+^T-cells from paired CSF and blood samples of untreated PwMS and control donors with idiopathic intracranial hypertension (IIH)(Kendirli et al., 2023). To avoid any bias, we adopted the reported bioinformatic strategy and regenerated the same UMAPs (Figure 2A). Considering all differentially expressed genes (DEGs) in the 13 clusters (sTable 1), as well as selected subset-associated marker genes (sFigure 4A-C), we annotated clusters containing cTFH (Cl 1), T_CM_ (CL 2), cells with an type-1 IFN signature (Cl 3), Th17-cells (Cl 4), GzmK^lo^Th1-like cells (Cl 8), Th2-cells (Cl 10) CD4^+^CTL (Cl12) and FOXP3^+^Tregs (Cl13) Figure 2A). Clusters 5, 6 and 9 lacked a clear gene signature and were annotated as T_EM_A, B and C. Cluster 7 displayed the gene signature of EOMES^+^Tr1-like cells (Bonnal et al., 2021; De Simone et al., 2021; Gruarin et al., 2019), as evidenced by a highly significant Gene Set Enrichment Analysis (GSEA, Figure 2B). Indeed, several key EOMES^+^Tr1-like signature genes, namely EOMES (Gruarin et al., 2019) (Figure 2C), CHI3L2 (Bonnal et al., 2021), CCL4, CCR5 and PDCD1 (PD1) (Alfen et al., 2018) were selectively expressed in this cluster (sFigure 4C). Moreover, they expressed the highest levels of GzmK, low levels of IL-7R and lacked CCR6 (Figure 2A, sFigure 4C). Also cells in Cluster 11 expressed GzmK and EOMES, but in contrast to Tr1-like cells they expressed CCR6 and high levels of IL-7R (Figure 2D), suggesting that this cluster contained pro-inflammatory CCR6^+^GzmK^+^ cells (sFigure 2). The CCR6^+^GzmK^+^ cluster was separated in the UMAP from EOMES^+^Tr1-like cells by a GzmK^lo^Th1-like cluster (Cl 8, Figure 2A), suggesting that they had largely diverse gene expression patterns. Indeed, cells in the CCR6^+^GzmK^+^ cluster expressed lower levels of EOMES^+^Tr1-like signature genes but very high levels of genes characteristic for pro-inflammatory Th1/17-cells (Figure 2E, sFigure 4C, sTable 2). Notably, the EOMES^+^Tr1-like cluster in the CSF of PwMS expressed the highest levels of EOMES^+^Tr1-like signature genes (Figure 2E). They had also an altered expression of genes regulating cholesterol metabolism, but this was a general feature of CSF-infiltrating CD4^+^T-cells (sTable 3, sFigure 5A)(Hrastelj et al., 2021).

**Figure 2:**
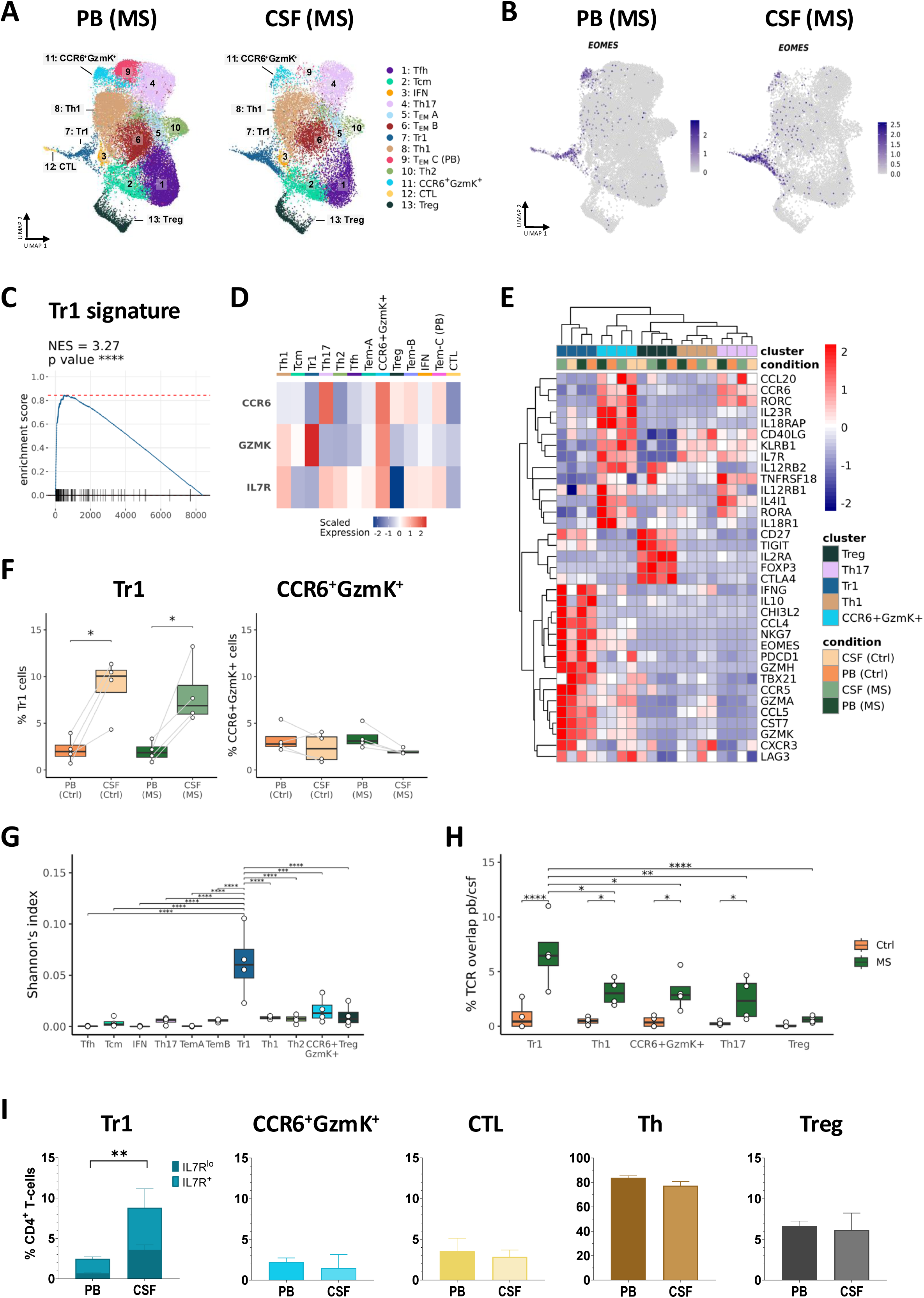
EOMES^+^Tr1-like cells generate a distinct cluster among single-sequenced CD4^+^T-cells and are enriched and clonally expanded in the cerebrospinal fluid of PwMS. **A.** UMAPs of 13 CD4^+^T-cell clusters in the blood and the CSF of 4 PwMS. **B.** UMAPs showing EOMES expression in the PB and the CSF of PwMS. **C.** Gene Set Enrichment Analysis of reference EOMES^+^Tr1-like signature genes of Cluster 7 (Overlay of all conditions). The normalized enrichment score (NES) and its p-value are reported in the panel. **D.** Heatmap of CCR6, GzmK and IL-7R expression in the 13 clusters (Overlay of all conditions). **E.** Heatmap of selected differentially expressed genes of EOMES^+^Tr1-like cells, CCR6^+^GzmK^+^ cells and FOXP3^+^Tregs. Th17 and Th1 clusters (Cl 4, 8) are shown as controls. Relative average expression levels in the CSF and in the blood of PwMS and control donors are shown. **F.** Frequencies of Cluster 7 (EOMES^+^Tr1-like) and 11 (CCR6^+^GzmK^+^) in the blood and the CSF of PwMs and control donors. Statistical significance was calculated with a paired t-test. **G.** Normalized Shannon’s index showing the rate of clonal expansions per cluster in the CSF of PwMS. Higher values reflect lower diversity. Statistical significance was calculated with one-way ANOVA. **H.** TCR clonotype sharings among cells from T-cell clusters analysed in the Heatmap (E) in the blood and the CSF in PwMS and control donors. Statistical significance was calculated with two-way ANOVA. **I.** Frequencies of the indicated CD4^+^T-cell subsets in peripheral blood (PB, n=17) and the cerebrospinal fluid (CSF, n=5) of treatment-naïve PwMS were assessed by flow cytometry. Statistical significance was calculated with a t-test.

Importantly, exclusively the EOMES^+^Tr1-like cluster was strongly, selectively and significantly increased in the CSF as compared to the blood (Figure 2F, sFigure 5B). Conversely, the CCR6^+^GzmK^+^ cluster was not enriched in the CSF, and CD4^+^CTL (Cl 12) were even largely excluded. TCR clonotype analysis unveiled further that only Tr1-like cells were clonally expanded in the CSF of PwMS (Figure 2G). Conversely, no clonal expansions were detectable in the CSF of control patients, with the caveat of the low cell numbers that are characteristic for uninflamed CSF. Interestingly, in the blood of PwMS and in control patients only cytotoxic clusters, i.e. CD4^+^CTL, CCR6^+^GzmK^+^ and Tr1-like cells, contained expanded clones (sFigure 5C, D). Moreover, we observed a significantly higher clonotype sharing between the Tr1 clusters from the blood and the CSF in PwMS as compared to other clusters (Figure 2H), consistent with an efficient recruitment of Tr1-cells from the blood to the CSF in PwMS.

Finally, to corroborate the enrichment of EOMES^+^Tr1-like cells in the CSF, we analysed CD4^+^T-cell subsets also from the blood and the CSF in an independent cohort of treatment-naive RR-PwMS (Table 1) by flow cytometry. EOMES^+^Tr1-like cells were confirmed to be highly enriched in the CSF (Figure 2I). Other subsets were not increased. Finally, all CD4^+^T-cell subsets expressed significantly higher levels of CXCR3 in the CSF as expected (Geginat et al., 2017; Kendirli et al., 2023), and virtually all Tr1-like cells in the CSF were CXCR3^+^ (sFigure 5E).

Collectively, these results show that EOMES^+^Tr1-like cells can be identified in an unbiased manner according to their unique gene expression profile and are strongly enriched and clonally expanded in the CSF of PwMS.

### EOMES^+^Tr1-like cells are the major IL-10 producing T-cell subset in the CSF of PwMS

Some *ex vivo* sequenced T-cells (Kendirli et al., 2023) expressed IL-10 mRNA. They belonged mostly to the EOMES^+^Tr1-like cluster (Figure 3A). Moreover, most IL-10^+^ cells co-expressed GzmK, but not FOXP3 and they were predominantly IL-7R^lo^ (sFigure 6A), suggesting that they represent *in vivo* activated EOMES^+^Tr1-like cells. Consistently, some IL-7R^−^Tr1-like cells expressed also IFN-γ. In contrast to IL-10^+^T-cells, IFN-γ^+^ and also IL-2^+^ cells were however present in several clusters and most co-expressed IL-7R (sFigure 6A).

**Figure 3:**
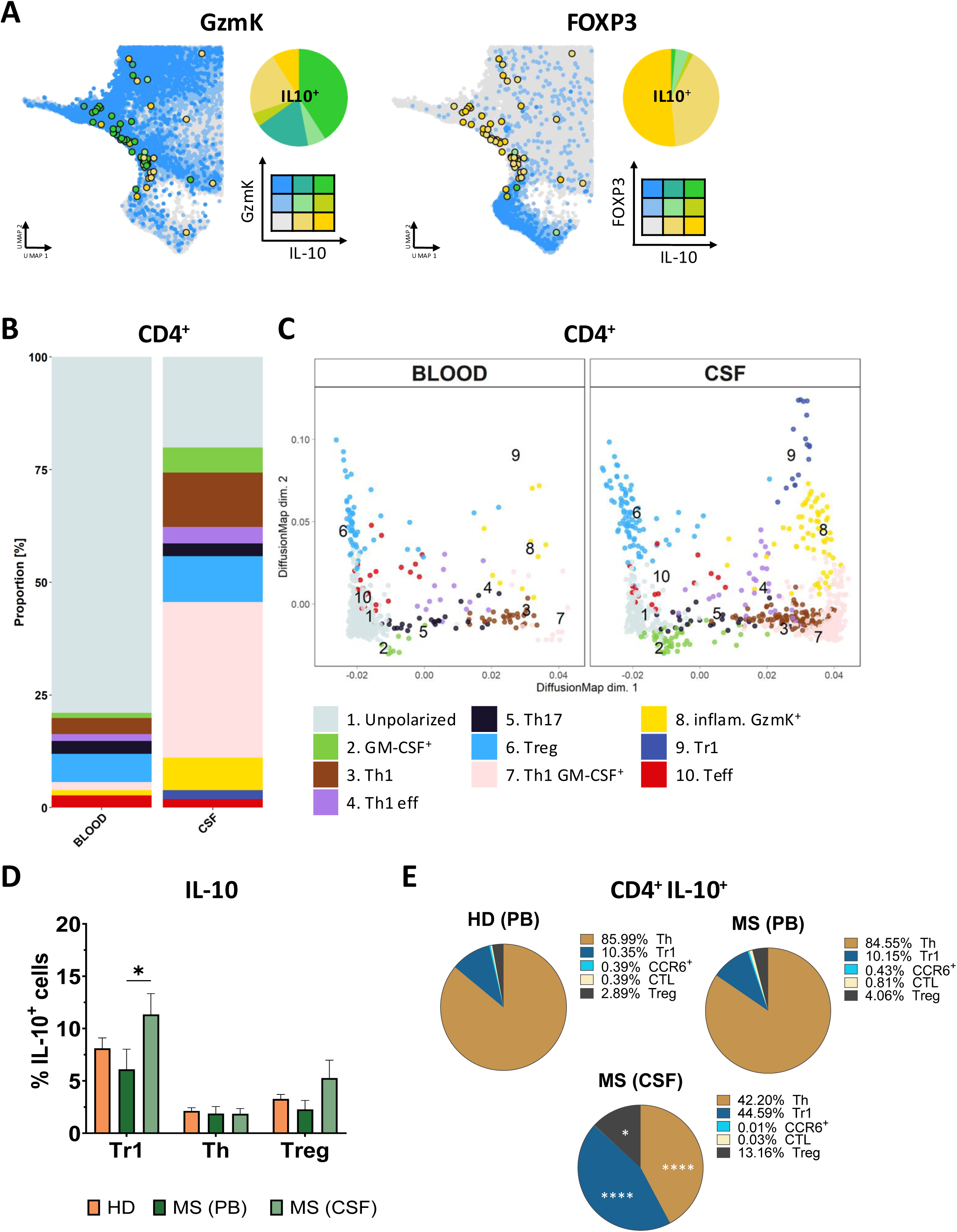
EOMES^+^Tr1-like cells produce high levels of IL-10 in the CSF of PwMS. **A.** UMAPS showing scaled gene co-expression of IL-10 with GzmK (left) or FOXP3 (right). The colour code and the relative scaled co-expression levels are reported in the panel. **B-E.** T-cells in PB and the CSF of untreated PwMS or HDs were activated with PMA and Ionomycin and analyzed for the expression of cytokines and differentiation markers. **B/C.** Clusters were identified and indicated by numbers, colors and names as indicated in the panel**. B.** Relative contributions of the 10 identified clusters to CD4^+^T-cells in PB and CSF **C.** Diffusion map analysis of the same clusters in PB and CSF predicting differentiation relationships in pseudotime. **D/E.** Analysis of IL-10 producing capacities of CD4^+^T-cell subsets in the blood of HDs (n=19) and in the blood and the CSF of PwMS (n=4). **D.** IL-10 production by the indicated CD4^+^T-cell subsets. Statistical significance was calculated by 2-way ANOVA **E.** Pie charts illustrating the relative contributions of the indicated T-cell subsets to the pool of IL-10-producing CD4^+^T-cells. Color code and mean percentages are reported in the panel. Statistical significance between PB and CSF of PwMS was calculated with a t-test.

We next analysed cytokine producing capacities of CD4^+^T-cells in the CSF and in peripheral blood of PwMS following brief stimulation with PMA and Ionomycin *ex vivo* by flow cytometry. Unsupervised clustering identified ten T-cell clusters, including a single IL-10^+^ cluster (Cl 9). The IL-10^+^ cluster co-expressed EOMES, GzmK and IFN-γ and corresponded thus to EOMES^+^Tr1-like cells (sFigure 6B). In the CSF there was a strong enrichment for clusters producing effector cytokines (Figure 3B, Cl 2-4 and 7-9). Cells in the effector cluster 8 contained pro-inflammatory EOMES^+^GzmK^+^cells (“inflam. GzmK^+^”) that expressed CD40L, IFN-γ and GM-CSF (Codarri et al., 2011; Pulvirenti et al., 2024). In the blood most CD4^+^T-cells expressed instead exclusively CD40L (Figure 3B, Cl 1, “unpolarized”). We next investigated possible differentiation pathways of the identified T-cell clusters by DiffusionMap analysis (Figure 3C, sFigure 6C): Dimension 1 separated clusters according to effector cytokines into a non-polarized and an effector group. Dimension 2 separated clusters furthermore according to CD40L expression into helper and regulatory/effector cells. IL-10^+^Tr1-like cells were positioned on top of the pro-inflammatory GzmK^+^ cluster, consistent with a precursor-effector relationship (Geginat et al., 2023; Pulvirenti et al., 2024).

Notably, EOMES^+^Tr1-like cells contained the highest percentages of IL-10^+^ cells following brief polyclonal stimulation of all CD4^+^T-cell subsets in both HDs and in PwMS (Figure 3D). In contrast to previous reports (Astier et al., 2006; Sumida et al., 2018), IL-10 producing capacities of CD4^+^T-cell subsets were not significantly reduced in the blood of PwMS, but they were significantly increased in EOMES^+^Tr1-like cells in the CSF. Notably, EOMES^+^Tr1-like cells could produce high amounts of IL-10 in the CSF of both RR- and P-PwMS (sFigure 6D). A completely different pattern was as expected observed for IFN-γ (sFigure 6E). Thus, IFN-γ producing capacities were in general very high in all cytotoxic subsets. Conversely, IFN-γ producing capacities were low among Th-cells and Tregs in the blood, but they were strongly increased in the CSF (Dominguez-Villar et al., 2011).

Although only a small fraction of Th-cells produced IL-10, they represent the large majority of CD4^+^T-cells and constituted consequently also the majority of IL-10 producing T-cells in the blood (Figure 3E). Conversely, in the CSF of PwMS the major IL-10 producing T-cell subset were EOMES^+^Tr1-like cells, and also the contribution of Tregs was significantly increased.

In conclusion, regulatory T-cells-predominantly EOMES^+^Tr1-like cells-could produce high amounts of IL-10 in PwMS, in particular in the CSF.

### Regulatory T-cells in PwMS respond poorly to myelin antigens but are activated by EBV to produce IFN-γ and IL-10

Antigenic activation induces T-cell proliferation and IL-10 production. We therefore investigated if regulatory T-cells in MS patients reacted with a panel of self- and viral antigens that we showed previously activate CSF-infiltrating CD4^+^T-cells in PwMS (Paroni et al., 2017). We modified an established assay to detect rare antigen-specific cells upon *ex vivo* stimulation with antigenic peptides according to activation marker induction (Bacher et al., 2013; Saggau et al., 2024), combining it with intracellular IL-10 and IFN-γ staining (sFigure 7). The assay allows to monitor *ex vivo* frequencies and qualities of antigen-specific T-cells even in a small amount of blood, which is often a limiting factor. Importantly, even rare antigen-specific T-cells (<1:10.000) could be detected with high reproducibility (sFigure 7F). Nevertheless, T-cell responses to the myelin-derived self-antigens MOG/MBP were hardly detectable (Figure 4A, sFigure 7E). Only CD8^+^T-cells and Th-cells showed statistically significant responses in respectively untreated and Natalizumab-treated PwMS as compared to the no peptide control (sFigure 8A). Importantly, EOMES^+^Tr1-like cells largely failed to respond at all to myelin antigens, including MOG, MBP and PLP (Figure 4A and sFigure 8A, B). In contrast, we detected significant IFN-γ responses to EBV, i.e. to a commercially available pool of immunodominant peptides, of EOMES^+^Tr1-like cells in HD and PwMs (Figure 4A, sFigure 8A). The response in HDs was low, but FACS-purified Tr1-like cells from HD proliferated with EBV, maintaining GzmK and EOMES expression (Bonnal et al., 2021) (sFigure 7G), confirming thus that EOMES^+^Tr1-like cells respond to EBV-derived antigens with a different assay. Notably, however, proliferation-based assays underestimate antigen-specific responses of anergic regulatory and exhausted/senescent effector T-cells (Geginat et al., 2003; Haringer et al., 2009). Importantly, Th-cells and in particular EOMES^+^Tr1-like cells responded strongly to EBV in Natalizumab-treated PwMS. Conversely, CD4^+^CTL and CCR6^+^GzmK^+^ subsets responded poorly or not at all. Notably, CD8^+^T-cells showed a strong response to EBV in HDs that was further increased in PwMS, independently of Natalizumab treatment (sFigure 8A). T-cell responses to JCV, a persistent virus that may cause progressive multifocal leukoencephalopathy (PML) in Natalizumab-treated PwMS, showed a completely different pattern. Thus, JCV activated predominantly Th-cells and CD4^+^CTL, but only weakly CD8^+^T-cells and EOMES^+^Tr1-like cells. Moreover, JCV-specific CD4^+^T-cell responses were overall reduced in treatment-naïve PwMS as compared to HDs and to Natalizumab-treated PwMS (Figure 4A).

**Figure 4:**
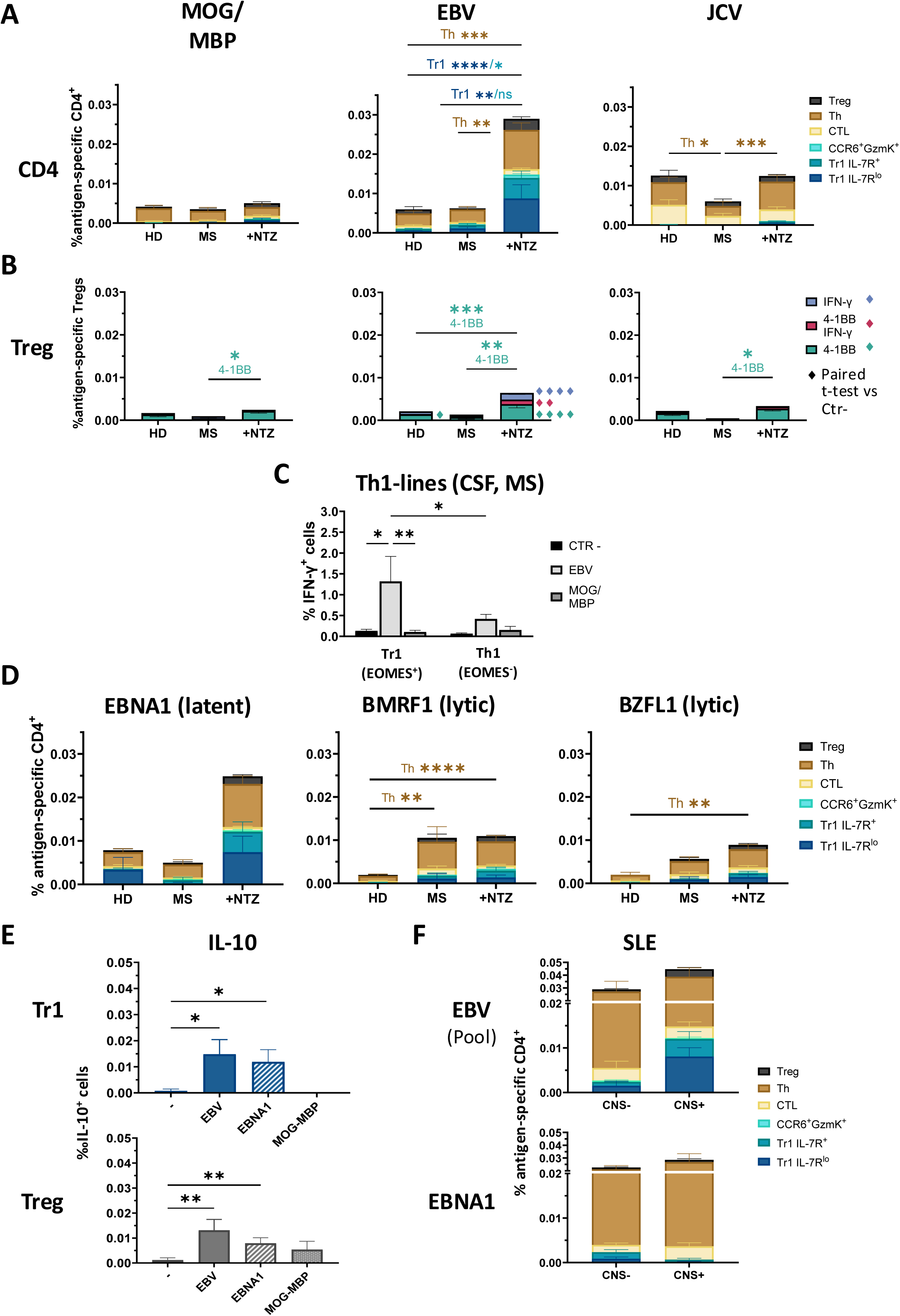
EOMES^+^Tr1-like cells are selectively activated by the EBV antigen EBNA1 in PwMS. **A.** Peripheral blood mononuclear cells were stimulated with peptide pools derived from MOG plus MBP, EBV (immunodominant mix) or JCV (VP1 plus VP2). T-cell subsets were tracked according to differentiation markers, and antigen-specific cells identified by concomitant CD69 upregulation and IFN-γ production (sFigure 7). Frequencies of antigen-specific cells in the indicated subsets are shown as stacked histograms among total CD4^+^T-cells, according to the indicated color code. Antigen-specific responses were analysed in HDs (MOG/MBP n=12, EBV n=24 and JCV n=10), treatment-naive (n=9, 10 and 8) and Natalizumab-treated PwMS (n=20, 27 and 16). **B.** Antigen-specific FOXP3^+^Tregs were tracked according to 4-1BB and/or IFN-γ expression among CD69^+^Tregs (HD n=10-24, untreated RR-PwMS n=2-5, Natalizumab-treated RR-PwMS n=10-15). **C.** CSF-derived CXCR3^+^CCR6^−^“Th1”-cell lines from 5 treatment-naive MS patients were stimulated with EBV or MOG/MBP-derived peptides, and IFN-γ production in EOMES^+^ and EOMES^−^ cells analysed. **D.** Antigen-specific IFN-γ responses to overlapping peptide pools derived from EBNA1, BMRF1 and BZLF1 (HD: n=6-10; untreated RR-PwMS: n=2-6; Natalizumab-treated RR-PwMS: n=13-26). **E.** IL-10 production by EOMES^+^Tr1-like cells and FOXP3^+^Tregs in response to EBV (immunodominant pool), EBNA1 (n=16) or MOG plus MBP (n=5) in Natalizumab-treated PwMS. **F.** Antigen-specific IFN-γ responses in SLE patients stratified according to CNS involvement (CNS-: n=11, CNS+: n=6) to EBV (upper panel) or EBNA1 (lower panel).

**Table 2:**
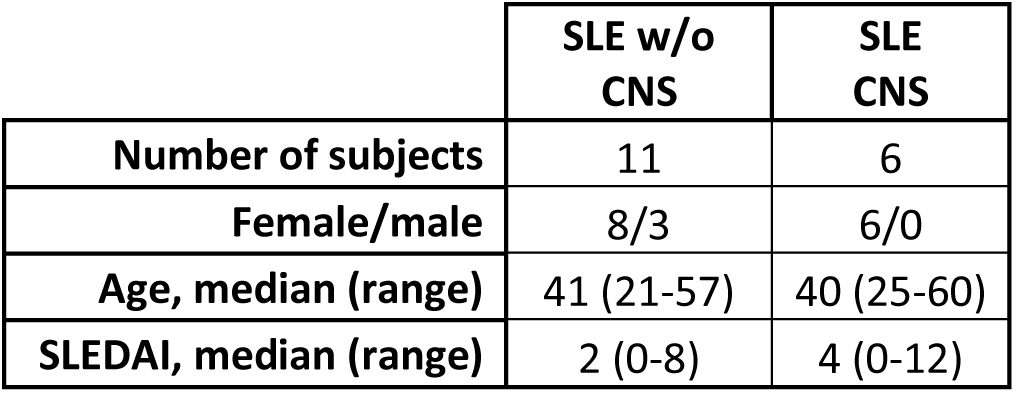
Characteristics of SLE patients with and without CNS involvement.

Interestingly, we detected also some IFN-γ production by FOXP3^+^Tregs in response to EBV (Figure 4A). Since only a minority of FOXP3^+^Tregs can produce IFN-γ (sFigure 6E), we analysed antigen-specific Treg responses also according to 4-1BB induction (Figure 4B), a well-established read-out of TCR-induced Treg activation (Schoenbrunn et al., 2012). The antigen-specific responses of FOXP3^+^Treg were overall similar to EOMES^+^Tr1-like cells. Thus, they showed a weak EBV-specific response in HDs but responded strongly and significantly in Natalizumab-treated PwMS. Responses of FOXP3^+^Tregs to myelin-derived antigens and JCV were in contrast low.

The strong increase of EBV-specific regulatory T-cells in Natalizumab-treated patients suggested that they could migrate to the CNS. Consistently, we showed previously that CSF-derived CCR6^−^CXCR3^+”^Th1”-cell lines of PwMS contained EBV-specific cells (Geginat et al., 2017; Paroni et al., 2017). EOMES expression is stably maintained in CD4^+^T-cell lines (sFigure 7G)(Bonnal et al., 2021), and here we found that a relevant fraction of CSF-derived “Th1”-cell lines expressed EOMES (sFigure 8C), suggesting that they contained EOMES^+^Tr1-like cells in addition to conventional EOMES^−^Th1-cells. These Eomes^+^Tr1-like cells showed a robust IFN-γ response to EBV-derived peptides (Figure 4C), suggesting that EOMES^+^Tr1-like cells in the CSF respond to EBV. Also EOMES^−^Th1-cells responded as expected to EBV, but significantly weaker.

EBV establishes a persistent, predominantly latent infection in B-cells, but can switch to lytic infection that leads to the expression of several immunogenic proteins (Murata et al., 2021). In Natalizumab-treated PwMS, EOMES^+^Tr1-like cells were strongly activated by a peptide pool of EBNA1 (Figure 4D, sFigure 9A), the most consistently expressed viral protein in latently infected B-cells. Notably, in a patient that donated enough blood to sort EOMES^+^Tr1-like cells, we observed also a strong proliferative response to EBNA1 (sFigure 9B), confirming thus EBNA1 specificity of Tr1-cells with an independent assay. In marked contrast, EOMES^+^Tr1-like cells responded poorly to two immunogenic lytic antigens (Figure 4D). Interestingly, Th-cells, CD8^+^T-cells and FOXP3^+^Tregs responded in contrast also to lytic EBV antigens, in particular in Natalizumab-treated patients (sFigure 9A).

We then investigated if EBV-specific T-cells could produce IL-10. Indeed, we were able to detect significant IL-10 production by antigen-specific EOMES^+^Tr1-like cells in response to the immunodominant EBV peptide pool and to EBNA1, whereas IL-10 was undetectable with myelin-derived peptides (Figure 4E). Similar results were obtained with FOXP3^+^Tregs. Conversely, antigen-specific IL-10 responses of Th-cells were variable and did not reach statistical significance (sFigure 9C).

To understand if EBNA1-specific regulatory T-cells were a unique feature of MS, we finally analyzed patients with systemic lupus erythematosus (SLE), a systemic autoimmune disease that is also associated with EBV and may affect the CNS. Interestingly, EOMES^+^Tr1-like cells and FOXP3^+^Tregs showed increased responses to EBV in SLE patients with CNS involvement as compared to SLE patients without CNS involvement (Figure 4F, sFigure 10A, B). Nevertheless, they failed to respond to EBNA1.

In conclusion, regulatory T-cells, particularly EOMES^+^Tr1-like cells, are activated by the latency-associated EBV-derived antigen EBNA1 in PwMS to produce IFN-γ and IL-10, but they fail to respond to myelin-derived autoantigens.

### EBNA1-specific EOMES^+^Tr1-like cells are associated with anti-EBNA1 IgG in PwMS and disappear following therapeutic B-cell depletion

We next asked if EBV-specific regulatory T-cells cells were associated with the major genetic risk factor of MS, the MHC class-II allele HLA-DRB1*15:01, which present antigenic peptides to activate CD4^+^T-cells (Tschochner et al., 2016). Individuals carrying this risk allele were enriched among the analysed Natalizumab-treated PwMS (Table 3). IFN-γ responses of EOMES^+^Tr1-like cells in HLA-DRB1*15:01^+^ PwMS to EBV-derived peptides were overall lower, but they were clearly detectable and not significantly reduced (Figure 5A). EBV-specific IFN-γ responses of Th-cells and Tregs were also similar, and also EBV-induced IL-10 production by EOMES^+^Tr1-like cells and FOXP3^+^Tregs was not significantly altered (sFigure 11A).

**Figure 5:**
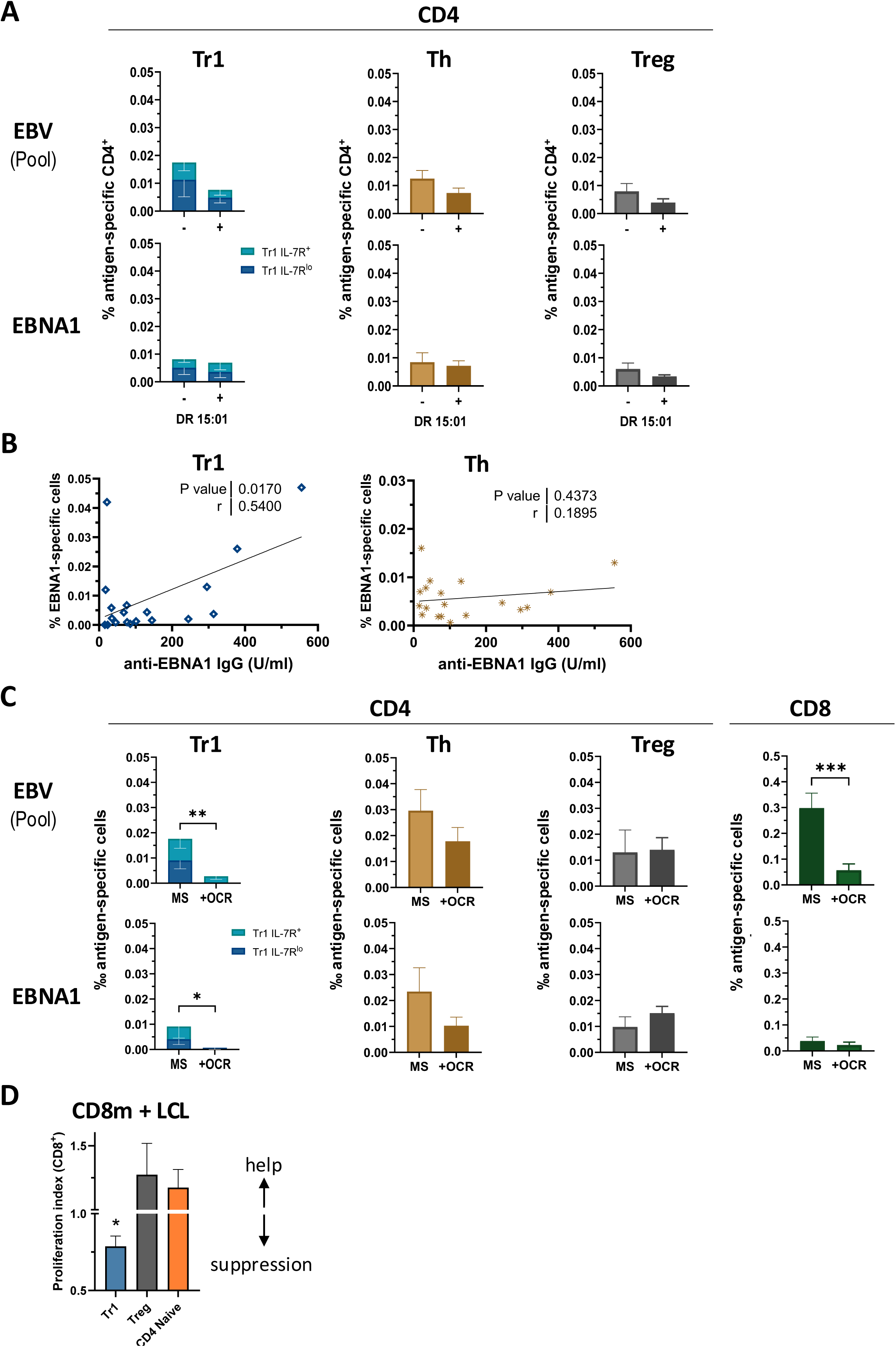
EBNA1-specifc EOMES^+^Tr1-like cells are associated with anti-EBNA1 IgG levels and disappear in Ocrelizumab-treated patients. **A.** Natalizumab-treated PwMS were HLA-haplotyped (Table 3) and the frequencies of EBV- or EBNA1-specific EOMES^+^Tr1-like cells, Th-cells and FOXP3^+^Tregs (Figure 4) stratified according to the expression of HLA-DRB1*15:01.**B.** EBNA1-specific EOMES^+^Tr1-like cells or Th-cells in Natalizumab-treated patients (n=11-19) were plotted against the serum levels of anti-EBNA1 IgG antibodies. R and p-values are reported on the right of the panels. **C.** EBV-specific T-cell response in untreated (n (RR, P)=10, 3) and Ocrelizumab-treated PwMS (n(RR, P)=6, 11)). Shown are responses of the indicated EBV-reactive CD4^+^T-cell subsets and of total CD8^+^T-cells to EBV (immunodominant pool, upper panels) and to EBNA1 (lower panels). **A/C.** Statistical significances were calculated with an unpaired t-test. **D.** Suppression of CD8^+^ memory T-cell proliferation induced by EBV-infected B-cells (LCL) and SEB by regulatory T-cell subsets. The proliferation index of CD8^+^ responder cells alone was set to 1; values of <1 indicate suppression, values >1 help. Naive CD4^+^T-cells are shown as negative control. Statistics were calculated with a paired t-test.

**Table 3:** HLA-DRB1 polymorphisms of Natalizumab-treated PwMS.

| Patient | 1 | 2 | 3 | 4 | 5 | 6 | 7 | 8 | 9 | 10 | 11 | 12 | 13 | 14 | 15 | 16 | 17 | 18 | 19 | 20 | 21 | 22 |
| --- | --- | --- | --- | --- | --- | --- | --- | --- | --- | --- | --- | --- | --- | --- | --- | --- | --- | --- | --- | --- | --- | --- |
| **HLA-DRB1\*** | 11:04<br>13:02 | 03:01<br>11:04 | 01:01<br>08:01 | 11:01<br>13:01 | 11:01<br>12:01 | 13:01<br>15:01 | 11:04<br>15:02 | 11:01<br>11:01 | 11:01<br>11:04 | 15:01<br>16:01 | 07:01<br>15:01 | 13:05<br>15:01 | 11:04<br>15:01 | 15:01<br>16:01 | 13:01<br>13:02 | 11:01<br>13:03 | 04:02<br>11:01 | 04:01<br>15:01 | 11:01<br>16:01 | 01:02<br>11:04 | 11:01<br>16:01 | 11:01<br>15:01 |

Anti-EBNA1 IgG antibody levels represent a strong independent environmental risk factor of MS (Bjornevik et al., 2023). Consistently, anti-EBNA1 IgG were significantly increased in both untreated and Natalizumab-treated PwMS compared to HDs (sFigure 11B). EBNA1-specific, IFN-γ-producing CD4^+^T-cells were weakly associated (p-value: 0.0513) with anti-EBNA1 IgG serum levels in Natalizumab-treated PwMS, and a weak tendency was also observed for CD8^+^T-cells (sFigure 11C). Intriguingly, a stronger and highly significant association was found for EBNA1-specific EOMES^+^Tr1-like cells (Figure 5B). Conversely, no association was found for EBNA1-specific FOXP3^+^Tregs (sFigure 11D). Moreover, there was also no association with EBNA1-specific Th-cells, irrespectively if they were identified according to IFN-γ (Figure 5B) or CD40L expression (sFigure 11E). Notably, EBNA1-activated EOMES^+^ and FOPXP3^+^ regulatory T-cells largely failed in contrast to Th-cells to upregulate CD40L (Facciotti et al., 2016; Schoenbrunn et al., 2012) (sFigure 11F), which is an absolute requirement for IgG induction (Gordon, 1995).

We then asked if EBV-specific regulatory T-cells were affected by Ocrelizumab, a therapeutic antibody depleting B-cells that could present EBV-derived peptides to T-cells. Ocrelizumab-treated patients (Table 1) had strongly and significantly reduced CD8^+^T-cell responses to EBV (Abbadessa et al., 2024), consistent with ongoing T-cell activation by EBV antigen-presenting B-cells in PwMS. Intriguingly, among CD4^+^T-cells only EBV-specific EOMES^+^Tr1-like cells were significantly reduced, and they were in most cases undetectable (Figure 5C). Conversely, EBV-specific FOXP3^+^Tregs were not affected and EBV-specific Th-cells were only moderately reduced.

Since both EOMES^+^Tr1-cells and FOXP3^+^T-cells could suppress CD8^+^T-cell proliferation in standard suppression assays (Bonnal et al., 2021; Pulvirenti et al., 2024), we investigated here if they could also suppress CD8^+^ memory T-cell activation induced by EBV-infected B-cells (LCL, Figure 5D, sFigure 12). EOMES^+^Tr1-like cells inhibited LCL-induced CD8^+^T-cell proliferation in a significant manner (Figure 5D). Surprisingly, FOXP3^+^Tregs failed to suppress and behaved overall similar to naïve CD4^+^ control cells. Thus, they provided even help in some experiments (Figure 5D, sFigure 12).

In conclusion, EBNA1-specific EOMES^+^Tr1-like cells are present in PwMS expressing the HLA risk allele, were associated with anti-EBNA1 IgG, disappeared following therapeutic B-cell depletion and suppressed the activation of CD8^+^ memory T-cells by EBV-infected B-cells.

### EOMES^+^Tr1-like cells produce IL-10 in brain lesions of PwMS

Finally, to assess if EOMES^+^Tr1-like cells were present in the CNS of PwMS, postmortem brain sections of P-MS cases were stained for CD4, EOMES and IL-10 (Figure 6, sFigure 13). EOMES^+^IL-10^+^CD4^+^ cells were identified in three different MS cases. They were found in the perivascular infiltrates of one active and one chronic active white matter lesion (Figure 6A, D-F) and in the inflamed meninges (Fig. 6H and insets). Cells co-expressing CD4 and EOMES, but not IL-10, were also detected (brown/red arrows in Figure 6A, H). They could represent either pro-inflammatory cells (Raveney et al., 2021) or EOMES^+^Tr1-like cells that were not activated to produce IL-10 at the analysed time point. EOMES was as expected also detectable in CD4^−^ cells (brown arrows in Figure 6A, H), and double staining of serial brain sections with anti-CD8 and anti-EOMES antibodies revealed the presence of EOMES^+^CD8^+^ cells in the same areas (Figure 6, inset in I), suggesting a colocalization of EOMES^+^Tr1-like cells and CD8^+^ memory/effector T-cells in the brain of PwMS.

**Figure 6:**
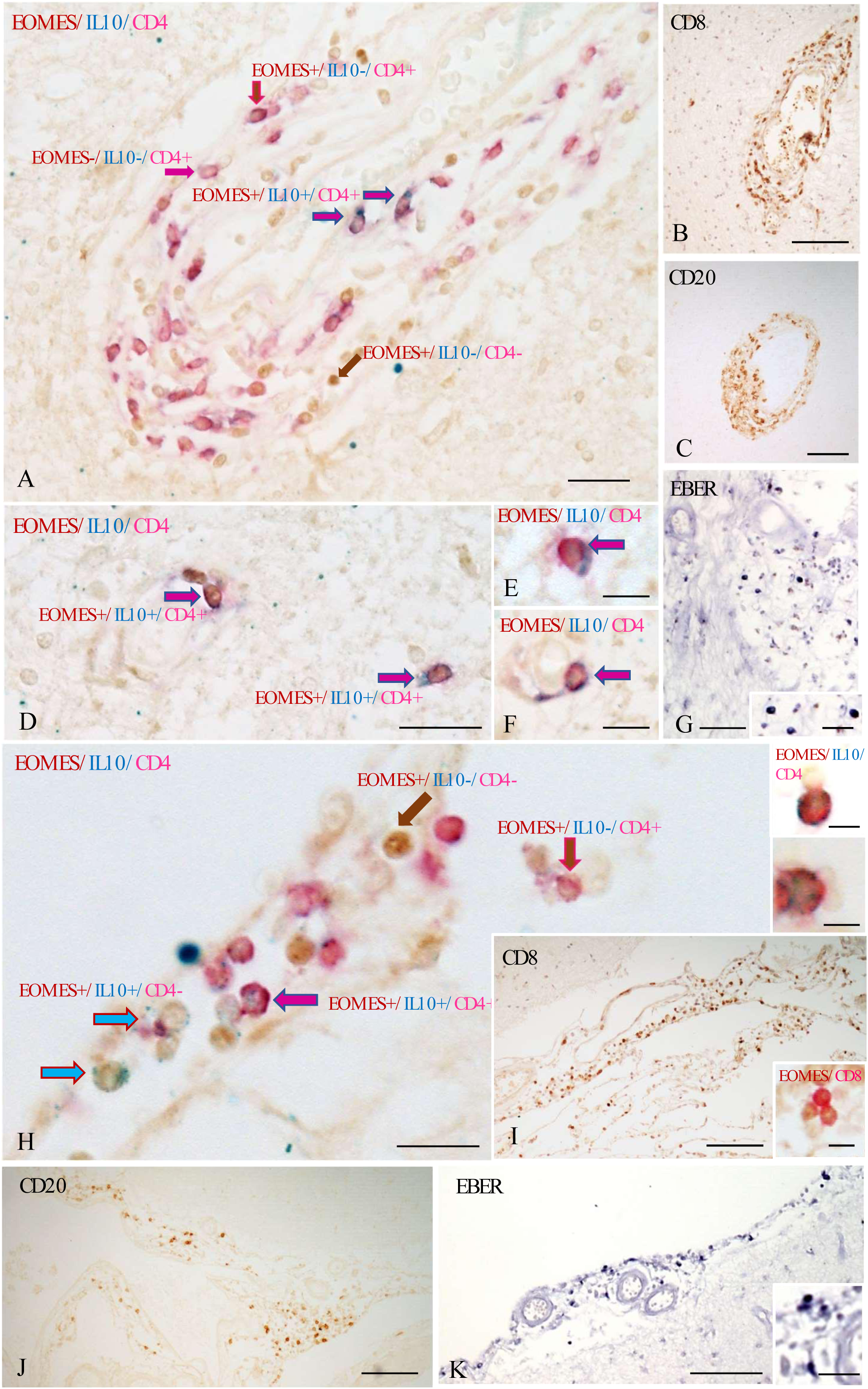
EOMES^+^ Tr1-like cells produce IL-10 in MS brain lesions. Triple immunostaining for EOMES (nuclear brown staining), IL-10 (blue) and CD4 (red) reveals presence of EOMES^+^IL-10^+^CD4^+^Tr1-like cells in perivascular immune infiltrates in one active WM lesion (blue/red arrows in A, D-F) and in the inflamed meninges (H) of one MS donor. In the same infiltrates, CD4^+^ cells expressing only EOMES (red/brown arrow in A and H), EOMES^−^ IL-10^−^CD4^+^ cells (red arrow in A) and EOMES^+^IL-10^−^CD4^−^ cells (brown arrow in A and H) were also detected. Panels E and F, inset in D and the left insets in H show four EOMES^+^IL-10^+^CD4^+^ cells at high power magnification. In the right inset in H, two EOMES^−^IL-10^+^ cells (arrows) one of which co-expresses CD4 (white arrow) are shown at high magnification. Immunostaining for CD8 (B and I) and double staining for CD8 and EOMES (inset in I), performed in serial sections, highlight the presence of numerous CD8^+^T-cells, some of which express EOMES, in the same immune infiltrates shown in A and H. Immunohistochemistry for CD20 (C and J) and *in situ* hybridization for the small non-coding EBV RNA (EBER) (G, K), performed in consecutive sections, reveal substantial B-cell infiltration and the presence of EBV-infected cells in the same brain areas. Bars: 100 µm in B, C, I-K; 50 µm in G; 20 µm in A, D, H and insets in G and K; 10 µm in E, F and inset in D, right inset in H, and inset in I; 5 µm in the insets in H.

CD4^+^ and CD4^−^ cells expressing IL-10, but not EOMES, were also detectable (Figure 6, insets in H), indicating that IL-10 production was not unique for EOMES^+^Tr1-like cells. We therefore counted cells in the brain sections to estimate the relative abundance of CNS-infiltrating EOMES^+^IL-10^+^CD4^+^Tr1-like cells. Approximately half of the IL-10^+^ cells expressed CD4. CD4^+^IL-10^+^EOMES^+^Tr1-like cells represented 6,3±3% (mean ± SD) of total CD4^+^T cells without any appreciable difference between meningeal and parenchymal inflammatory cell infiltrates. The percentages of CD4^+^ cells expressing either EOMES or IL-10 alone were more variable (EOMES^+^: 14±10%; IL-10^+^: 3.3±3.8%, respectively) and did again not differ significantly between meningeal and parenchymal immune infiltrates. Thus, EOMES^+^Tr1-like cells were a major source of IL-10 among CD4^+^T-cells also in these brain sections. Furthermore, approximately one third of CD4^+^EOMES^+^T-cells stained positive for IL-10, i.e. a higher fraction then those producing IL-10 upon stimulation with PMA and Ionomycin in the blood and the CSF (Figure 3d). No IL-10 was in contrast detected in intraparenchymal neural cells and microglia. Immunostaining for CD20 (Figure 6C, J), and *in situ* hybridization for the non-coding EBV-derived RNA EBER (Figure 6G, K), performed in consecutive brain sections, revealed in addition the presence of numerous EBV-infected B cells in the same immune infiltrates that contained EOMES^+^Tr1-like cells.

Overall, this analysis indicates that EOMES^+^Tr1-like cells are present in the MS brain, some in the vicinity of CD8^+^T-cells or EBV-infected B-cells and are locally activated to produce IL-10.

## Discussion

Eomes^+^CD4^+^T-cells contain both pro- and anti-inflammatory subsets, and have been proposed to play a pathogenic role in EAE (Stienne et al., 2016) and in progressive MS (Raveney et al., 2021). Our findings suggest a prominent role of anti-inflammatory EOMES^+^Tr1-like cells in MS. Thus, we documented here a dysregulated homeostasis selectively of EOMES^+^Tr1-like cells in MS that was related to their recruitment and activation in the CNS, where they were activated to produce IL-10. We further showed that these adaptive regulatory T-cells could be activated to produce IL-10 by the EBV antigen EBNA1, but not by myelin-derived antigens, suggesting that they inhibit anti-EBV immune surveillance but are insufficient to prevent CNS autoimmune pathology.

At variance with previous reports on defective IL-10 production by CD4^+^T-cells in MS (Astier et al., 2006; Sumida et al., 2018), and by our group in Inflammatory Bowel Diseases in the gut (Alfen et al., 2018), we found that EOMES^+^Tr1-like cells produced high amounts of IL-10 in the CSF and in the brain of PwMS. Thus, a decrease of IL-10 producing T-cells in the blood of PwMS is likely to reflect their recruitment to the CNS, but not a general defect of Tr1-cell differentiation or IL-10 production. IL-10 exerts predominantly anti-inflammatory functions in the CNS (Geginat et al., 2016). Nevertheless, also a role for T-cell derived IL-10 in sustaining pathogenic T cell survival and CNS inflammation in EAE was recently proposed (Yogev et al., 2022). In addition to IL-10, EOMES^+^Tr1-like cells produce high levels of IFN-γ, but IFN-γ has a neutral or even protective role in CNS inflammation (Dardalhon et al., 2008; Heinemann et al., 2014). EOMES^+^Tr1-like cells may however release cytotoxic molecules that could cause collateral CNS damage (Haile et al., 2015; Terrabuio et al., 2025).

IL-10 production and proliferation of adaptive regulatory T-cells are controlled by T-cell receptor stimulation. Therefore, the identification of the antigens that generate and activate these cells is a key understudied issue to understand their functions, in particular in MS (Paroni et al., 2017). In the EAE model, T-cells, including adaptive regulatory T-cells, are deliberately primed by myelin-derived self-antigens. Conversely, in PwMS regulatory T-cells largely failed to respond to three major CNS autoantigens. A failure to efficiently suppress myelin-targeted autoimmunity could explain the extensive CNS damage in MS. An unexpected key finding of this study is that regulatory T-cells in MS are activated by EBV, a persistent virus that plays an essential detrimental role in MS (Bjornevik et al., 2022). EOMES^+^Tr1-like cells responded selectively to EBNA1, the viral protein that is most consistently expressed in latently infected B-cells and therefore represents the virus’ Achilles heel (Murata et al., 2021). EBNA1 specific CD4^+^T-cells in PwMS were previously reported to represent Th1-cells (Lunemann et al., 2008), but since EOMES^+^Tr1-cells were identified later, these IFN-γ-producing CD4^+^T-cells contained probably both conventional Th1-cells and EOMES^+^Tr1-like cells. We showed that EBNA1-specific EOMES^+^Tr1-like cells were generated in MS, but not in SLE patients with CNS involvement and are thus likely to represent a MS-associated viral immune evasion strategy. Antibodies and T-cells that cross-react with EBNA1 and epitopes from self-proteins have been identified in PwMS (Geginat et al., 2017; Lunemann et al., 2008; Tschochner et al., 2016; Wucherpfennig and Strominger, 1995), suggesting that EBV infection might promote MS autoimmune pathology through molecular mimicry. EOMES^+^Tr1-like cells failed however to respond to myelin-derived antigens. Consistently, a previous study showed that EBV-specific Th1 clones that failed to cross-react with myelin antigens produced CCL4 rather than IL-2 (Lunemann et al., 2008), a pattern reminiscent of Tr1-cells (Haringer et al., 2009). We can of course not exclude that EOMES^+^Tr1-like are activated by other cross-reactive antigens (Tschochner et al., 2016), but molecular mimicry leading to the activation of regulatory T-cells in MS would be expected to have a paradoxical beneficial effect.

EOMES^+^Tr1-like cells lack B-cell helper functions and may suppress T-cell-dependent B-cell responses unless they up-regulate CD40L (Facciotti et al., 2016). Since also EBNA1-specific EOMES^+^Tr1-like cells in MS failed to up-regulate CD40L, it seems highly unlikely that they activate B-cells to produce anti-EBNA1 IgG antibodies. The association of EBNA1-specific IgG and EOMES^+^Tr1-like cells could rather reflect antigen presentation by EBNA1-specific B-cells, which can internalize EBNA1 via their BCRs with very high efficiency to present EBNA1-derived epitopes on MHC Class-II to CD4^+^T-cells (Lanzavecchia, 1985). Consistently, EBNA1-specific EOMES^+^Tr1-like cells disappeared from the CD4^+^T-cell compartment upon therapeutic B-cell depletion in PwMS, suggesting that they are continuously generated by EBV antigen-presenting B-cells. Anti-EBNA1 antibodies are produced by EBNA1-specific memory B-cells following differentiation to plasma cells and may promote MS by molecular mimicry. EBNA1-specific B-cells may in addition increase the risk of MS because they induce EBNA1-specific regulatory T-cells that favor viral immune escape.

CD8^+^T-cell responses to EBV are of broader magnitude in PwMS, and increased EBV reactivation and CD8^+^T-cell responses to lytic EBV antigens have been associated with active disease (Angelini et al., 2013). Interestingly, CD8^+^T-cells in PwMS recognise predominantly lytic antigens (Schneider-Hohendorf et al., 2022), suggesting that immune surveillance of latently infected B-cells may be inefficient. Intriguingly, immune-deficient mice reconstituted with human immune cells from HLA-DRB1*15:01-positive individuals had an inefficient control of EBV infection and mount dysregulated EBV-specific CD8^+^T-cell responses (Zdimerova et al., 2021), providing a link between the HLA risk haplotype and an inefficient EBV immune control. EBNA1-specific Tr1-like cells were not significantly reduced in PwMS carrying the HLA risk haplotype, indicating that EBNA1-derived epitopes that induce EOMES^+^Tr1-like cells are not necessarily HLA-DRB1*15:01-restricted. Finally, studies in post-mortem brain tissue from PwMS suggested that EBV, which can be detected in B-cell enriched inflammatory infiltrates in the CNS in a fraction of PwMS (Magliozzi et al., 2007), could lead to the recruitment and activation of EBV-specific CD8^+^T-cells (Magliozzi et al., 2007; Serafini et al., 2019). Our finding that EOMES^+^Tr1-like cells were present in the same CNS inflammatory infiltrates where EBV^+^B-cells and CD8^+^T-cells were detectable supports the concept that EOMES^+^Tr1-like cells could inhibit EBV-specific immune responses in the CNS. Indeed, EOMES^+^Tr1-like cells, but not FOXP3^+^Tregs, could suppress CD8^+^T-cell responses induced by EBV-infected B-cells *in vitro*. Thus, EBV-specific regulatory T-cells could inhibit CD8^+^T-cell-dependent immune surveillance of EBV (Geginat et al., 2017). This in turn could promote the generation of EBNA1-specific EOMES^+^Tr1-like cells due to persistent antigenic stimulation in a positive feed-forward loop. EBV-specific regulatory T-cells may thus be both a cause and a consequence of an inefficient anti-EBV immune control. CXCR3-dependent migration of EBV-infected B-cells to the CNS (Laderach et al., 2025; Saraste et al., 2016) may trigger the recruitment of virus-specific helper and regulatory T-cells (Angelini et al., 2013; Balashov et al., 1999; Lossius et al., 2014; Magliozzi et al., 2013; Serafini et al., 2007; Serafini et al., 2019; Serafini et al., 2010) and of auto-reactive Th1/17 bystander cells (Cao et al., 2015; Paroni et al., 2017; Wang et al., 2020), leading ultimately to CNS damage (Geginat et al., 2017). We thus propose that IL-10 produced by EBV-specific regulatory T-cells has a dual role in MS: It could initially prevent collateral CNS damage in prodromal MS that develops after EBV infections and is characterized by unsymptomatic CNS damage (Bjornevik et al., 2022). However, since it favors viral immune escape, it enhances also the risk of EBV-associated autoimmunity in the long run.

A lack of protection by regulatory T-cells is normally attributed to reduced Treg numbers or functions, namely IL-10 production in MS. Our findings suggest the novel concept that Treg dysfunction may also be a consequence of an aberrant antigen specificity, which promotes immune evasion of a disease-promoting pathogen but is inefficient to inhibit auto-reactive T-cells and immunopathology in the target tissue. A general implication of this study is further that the analysis of polyclonal adaptive Treg populations without the knowledge of their antigen specificities may be insufficient to understand their role in complex human pathologies.

## Materials and Methods

### Human samples

Patient samples were collected at the MS Centre of the *Fondazione Ca’ Granda, IRCCS Ospedale Policlinico*, Milan (“Policlinico”), upon written informed consent, after Institutional Ethical Committee approval (parere 708_2020). Patients were diagnosed according to the revised McDonald criteria(Hartung et al., 2019). We collected 6-8 ml of peripheral blood samples, by venous puncture, from 13 treatment-naïve patients affected by RR-MS (1-3d after an attack), 4 untreated P-MS patients, 22 Natalizumab-treated RR-MS patients and 17 Ocrelizumab-treated MS patients (containing both RR and P-MS). For the latter, blood was collected before the therapy session. 3-7 ml of CSF samples were obtained by lumbar puncture from 7 untreated patients at diagnosis. In 5 of these patients also paired blood samples were obtained. All patients whose CSF was analysed, were diagnosed as RR-MS and had not received any previous treatment. Demographic and clinical characteristics of MS patients and healthy donors (HD) enrolled in the study are shown in Table 1. As controls, 78 age- and sex-matched healthy donors were enrolled at the Policlinico upon informed consent.

### HLA genotyping

Genomic DNA was isolated from peripheral blood using Qiagen columns. DNA was isolated from frozen peripheral blood. DNA purification was performed with Qiagen EZ1 Advance XL automated instrument using EZ1 DSP DNA Blood Kit (QIAGEN). HLA-DRB1 typing was performed using Luminex based reverse SSO typing by using the LIFECODES HLA-DRB1 eRES SSO Typing SSO typing kit (Immucor).

### Cell isolation and stimulation

Peripheral blood mononuclear cells (PBMCs) were isolated from heparinized peripheral blood by Ficoll-Hypaque gradient within 24h from blood withdrawal. Cells from CSF were collected by centrifugation and analysed immediately. Plasma was collected and decomplemented or stored at −80°C until use. PBMC were incubated for 4h at 37°C in a 5% CO2 humidified atmosphere at a density of 2,5 x 10^6^ cell/mL in RPMI complete medium (2 mM glutamine, 1 mM sodium pyruvate, 1% non-essential amino acids, 1% penicillin/streptomycin) supplemented with 10% FCS in presence or absence of PMA (50 ng/mL) and Ionomycin (500 ng/mL). After 2h, Monensin (0.6 μl/ml,) was added and stimulated cells were stained for superficial and intracellular markers (listed in sTable 4). For antigen-specific stimulations cells were stimulated for 5 hours with peptide pools (EBV: Pool from 15 EBV proteins, or overlapping peptide pools from EBNA1, BZLF1, BMRF1; JCV: VP1 plus VP2; Myelin: MOG, MBP, PLP, Miltenyi) in medium without FCS that contained 5% autologous plasma. Incubation without peptides was analysed as negative control, and with 100 ng/ml of the super-antigen SEB as positive control. CD4^+^CXCR3^+^CCR6^−^“Th1” and CD4^+^CCR6^+^CXCR3^−^“Th17”-cells were sorted from the CSF of untreated PwMS, excluding IL-7R^lo^CD25^+^Tregs. Cell lines were generated as previously reported (Paroni et al., 2017) and restimulated with peptides as described above.

### Flow Cytometry

Cells were stained at the surface and intracellularly with fluorochrome-conjugated monoclonal antibodies and live/dead (L/D) exclusion dye. The different panels of antibodies (Ab) are represented in sTable 4. For surface staining, PBMCs were incubated for 20 min at 37°C, with the appropriate antibodies, including a L/D dye, diluted in Brilliant Staining Buffer (BD). To stain for intracellular molecules samples were fixed with the Fixation/Permeabilization working solution (eBioscience). For cytokine detection following PMA plus Ionomycin stimulation cells were fixed with 2% PFA. Fixed cells were permeabilized for 20 min at RT (eBioscience, Permeabilization Buffer). Cells were incubated with the antibodies at 4°C for 120 minutes. Samples were acquired on a BD FACSymphony and analysed with the BD FloJo Software. Antigen-specific CD4^+^ and CD8^+^T-cells were gated as CD69^+^IFN-γ^+^ unless otherwise indicated. Antigen-stimulated CD4^+^T-cell subsets cells were in some cases also analysed for IL-10 production or CD40L up-regulation. Tregs were gated as 4-1BB^+^CD69^+^ (Schoenbrunn et al., 2012). Notably, the markers used to identify T-cell subsets (CD4, CD8, IL-7R, CCR6, FOXP3, GzmK, GzmB (Cossarizza et al., 2021)) were stably expressed in the analysed time window (sFigure 4D).

### Suppression assay

LCL lines were cultured standard complete RPMI medium. CD8^+^CD45RA^−^ memory T-cells were sorted from PBMCs and stained with Cell Trace Violet Cell Proliferation Kit (INVITROGEN). CD4^+^T-cells were enriched using anti-CD4 magnetic microbeads (Miltenyi). Tr1 and Treg subsets as well as CD4^+^CD45RA^+^ naïve control cells were then isolated by FACS sorting. LCL and CD8 memory T-cells were seeded in 96 V shaped multiwell plates flasks in standard complete RPMI (CORNING) in the absence or presence of CD4^+^T-cell subsets and Staphylococcus enterotoxin B (SEB, 1 μg/mL) added. The ratio between LCL, CD8^+^ and CD4^+^ T-cells was 0.5:1:1 After 7 days of co-culture cells were collected and Cell Trace Dilution of CD8+ T-cells assessed. Analysis of FCS data was performed using BD FlowJo software (proliferation modeling tool).

### Anti-EBNA1 ELISA

ELISA to measure the levels of anti-EBNA1 IgG were performed using serum diluted 1:5 with a kit (Serion Diagnostics) following the manufacturer’s instructions.

### scRNAseq data

Processed single-cell RNA sequencing (scRNA-seq) gene expression data were obtained from the Gene Expression Omnibus (GEO) database under the accession number GSE232343 (Kendirli et al., 2023). The data were subsequently analyzed using the Seurat package (v.5.0.1) in R. The scRNA-seq dataset contained clusters defined in the original study. We utilized Seurat’s FindMarkers function with default parameters and Wilcoxon test to obtain Differentially Expressed Genes (DEGs) for each cluster. Clusters were labelled using the DEGs and additional information of known CD4^+^T-cell subset markers. To identify DEGs between conditions within the same CD4^+^T-cell subpopulation, we employed a pseudobulk approach. This method aggregates single-cell counts within each condition/patient to create bulk-like profiles, thereby improving the robustness of the differential expression analysis. Gene set enrichment analysis (GSEA) was performed using the fgsea() function from the fgsea package (v1.20.0). The reference Tr1 associated gene set for the analysis was obtained from our previous studies(Bonnal et al., 2021; De Simone et al., 2021). For TCR analysis, we selected cells with paired alpha and beta chain sequenced (69112 cells). TCR overlap was defined as the presence of identical paired alpha-beta chains in cells of the same cell-type across the two tissues. Clonal diversity within each cluster was assessed using the normalized Shannon’s diversity index. For a given cluster with *S* unique clonotypes and relative frequencies *p_i_*, the Shannon index was calculated as 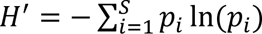. To account for differences in cluster size, was normalized by the maximum possible value ln(*S*). To better capture clonal expansion, we used the inverted form of the index: 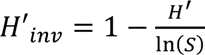, where higher values reflect reduced diversity and dominance by fewer clonotypes.

### Statistical analysis

Data were analysed with FlowJo software (Tree Star Inc, v. 10.8), while the statistical analyses were performed with Prism10 software (GraphPad). Student t-test were performed to compare two groups, a paired t-test for values derived from the same individuals, and an unpaired t-test to compare two different cohorts. One-way ANOVA was used to compare a single parameter in multiple groups, two-way ANOVA to compare multiple values in multiple groups. P-values <0.05 (*), <0.01 (**), < 0.001 (***), < 0.0001 (****) were regarded as statistically significant. Where indicated, borderline p-values (0.05-0.1) were reported in the panels.

### Tissue samples and immunostainings

Sections from 5 well-characterized post-mortem brain tissue blocks from 4 donors with progressive MS, provided by the UK Multiple Sclerosis Tissue Bank at Imperial College London (https://www.imperial.ac.uk/medicine/multiple-sclerosis-and-parkinsons-tissue-bank/) were analysed. The use of human tissue for research purposes was approved by the Ethics Committee of Istituto Superiore di Sanità (2023_0059846). Tissue sections used in this study contained substantial immune infiltrates with CD4+ and CD8+ T cells in the white matter (WM) and meninges. Triple immunostainings in bright field were performed with mouse anti-EOMES, rat anti-IL-10 and rabbit anti-CD4 purified monoclonal antibodies (see supplementary Methods). Negative control stainings were performed in consecutive sections using rat and mouse Ig isotype controls (ThermoFisher, sFigure 13). In serial sections immunostainigs for CD8, CD20 and EBER (Serafini et al., 2023) and double stainings with rabbit polyclonal anti-CD8 (Thermo Fisher Scientific) and anti-EOMES were also performed. Stained sections were analysed and images acquired with an Axioscope microscope (Carl Zeiss, Jena, Germany) equipped with AxioCam MRc5 camera and Axiovision SE 64 software. Cell counts were performed manually in the immune infiltrates containing CD4^+^ cells on the entire sections, both the WM and meninges, with a morphometric grid within the ocular lens, at 20x magnification. Data are expressed as mean ±SD values of the percentages of triple positive EOMES^+^IL10^+^CD4^+^ cells in the total CD4^+^T-cell population.

## Acknowledgments

This work was supported by FISM - Fondazione Italiana Sclerosi Multipla (2017/R/14) and co-financed with the ‘5 per mille’ public funding. This project has received also funding from the European Union’s Horizon Europe Research and Innovation Actions under grant no. 101137235 (BEHIND-MS). Views and opinions expressed are however those of the author(s) only and do not necessarily reflect those of the European Union nor the granting authority. Neither the European Union nor the granting authority can be held responsible for them. DG and ES are supported by the Italian Ministry of Health (Ricerca Corrente). We thank Manuel Albanese and Samuele Nortabartolo for critical reading and comments, and Francesca Aloisi for inspiring discussion and suggestions.

sTable 1: Differentially expressed genes in Cluster 1-13

sTable 2: Differentially expressed genes in EOMES^+^Tr1-like cells: CSF versus blood

sTable 3: Differentially expressed genes in EOMES^+^Tr1-like cells versus CCR6^+^GzmK^+^ cells

sTable 4: Flow Cytometry antibody panels

## Supplementary Figures

**sFigure 1: T-cell clusters in peripheral blood of HDs and PwMS identified by multiparametric flow cytometry.**

A. Histograms showing the expression of the 18 markers analysed to classify T-cells from peripheral blood of HDs and of PwMS in the 27 identified T-cell clusters. B. UMAP maps of the identified T-cell clusters showing the individual expression of the 15 most relevant markers. C. Unsupervised hierarchical clustering of the 12 CD4^+^T-cell clusters according to the expression of the indicated differentiation markers. Red indicates high and blue low expression. D. Relative contribution of the identified T-cell clusters to the T-cell compartments in PwMS and HDs

**sFigure 2: Gating strategy to identify pro- and anti-inflammatory EOMES^+^CD4^+^T-cell subsets**

A. CD4^+^T-cells were analysed according to the expression of the indicated markers to identify FOXP3^+^Tregs (left), GzmB^+^CTL, Th-cells (central panel) and GzmK^+^ subsets (right panel), namely IL-7R^+/lo^Tr1-like and CCR6^+^ cells. B. EOMES expression in the indicated T-cell subsets (n=9) in PwMS. C. Cytokine profiles of GzmK^+^CD4^+^T-cell subsets in PwMS

**sFigure 3: Frequencies and activation of CD4^+^T-cell subsets in PwMS and HDs**

A. Frequencies, CD69 and Ki67 expression of the indicated CD4^+^T-cell subsets in 35 HDs, 3 untreated P-PwMS, 11 treatment-naïve and 20 Natalizumab-treated RR-PwMS. B. CXCR3 surface expression on the indicated Ki67^+^T-cell subsets in the blood of treatment-naïve RR-PwMS (n=7-11).

**sFigure 4 Annotation of CD4^+^T-cell clusters.**

A. UMAPs of 13 antigen-experienced T-cell clusters in the blood and the CSF of 4 control donors (with idiopathic intracranial hypertension, IIH). B. Dot plot showing expression levels of the indicated signature genes of central memory, Tfh, Th1, Th2, Th17, CTL, EOMES^+^Tr1-like, Treg and IFN-activated cells in the 13 CD4^+^T-cell clusters C. Upper UMAPS: Individual signature genes of Tfh (Cl1, CXCR5), IFN activate cells (Cl3, IFIT3), Th2 (Cl 10, PTGDR2/CRTH2), CTL (Cl12, GzmB) and Treg (Cl13, FOXP3). Central UMAPS: Selective expression of EOMES^+^Tr1-like-associated signature genes (CHIL2, CCL4, CCR5 and PDCD1 (PD1) in Cluster 7. Lower UMAPS: Expression of GzmK and of the indicated Th1/17-associated genes in Cl 11(CCR6, RORC, IL-23R, IL-18RAP). All UMAPS show overlays of all conditions (Blood, CSF, MS, IIH).

**sFigure 5 Features of EOMES^+^Tr1-like cells in the CSF and the Blood.**

A. Heatmaps showing the average expression of genes upregulated in CSF-infiltrating versus circulating EOMES^+^Tr1-like cells involved in Cholesterol metabolism and selectively down-regulated gens in EOMES^+^Tr1-like cells (upper) and all other CD4^+^T-cells (lower panel) in the CSF and in the blood. B. Relative contributions of the identified clusters to total antigen-experienced CD4^+^T-cells in the blood and the CSF of PwMS and control Donors. C. Normalized Shannon’s index showing the rate of clonal expansion per cluster in the PB of PwMS, PB and CSF of Controls, respectively. Higher values reflect lower diversity. Statistical significance was calculated with one-way ANOVA. D. Bubble plot illustrating the clonal expansion of Treg, Tr1, CCR6⁺GzmK⁺, and CTL in the CSF and PB of PwMS. Bubble radius is proportional to the number of cells sharing the same TCR within each patient. Clones comprising more than two cells are annotated with their cell counts. E. CXCR3 surface expression in the indicated CD4^+^T-cell subsets in peripheral blood (PB, n=11) and in the CSF of PwMS (n=3). Statistical significance was calculated by One-way ANOVA.

**sFigure 6 Cytokine profiles of CD4^+^T-cell subsets.**

A. UMAPs showing the scaled co-expression of IL-10, IFN-γ and IL-2 with IL-7R. Color code and relative scaled co-expression levels are reported in the panel. B. Hierarchical clustering of the 10 identified CD4^+^ T-cell clusters according to the expression of cytokines and differentiation markers in the blood and CSF of active MS patients. C. Diffusion Maps showing the expression of 7 principal markers that characterize the cytokine producing T-cell clusters. D. IL-production by EOMES^+^Tr1-like cells in HDs (n=19) and RR-(n=2-3) or P-PwMS (PB n=2) in the blood or the CSF. E. Analysis of IFN-γ producing capacities of CD4^+^T-cell subsets in the blood of HDs (n=19) and in the blood and the CSF of PwMS (n=4).

**sFigure 7: Assessment of antigen-specific T-cell subsets *ex vivo*.**

A. TCR-induced T-cell responses identified by concomitant CD69 and IFN-γ induction in gated CD4^+^T-cells. B. TCR-induced responses of FOXP3^+^Tregs identified by concomitant CD69 and 4-1BB induction. C. TCR-induced upregulation of CD40L versus IL-10 on total CD4^+^T-cells. A-C. Shown are the negative (Ctr-, no antigen) and the positive (Ctr+, super-antigen SEB) control. D. Stable expression of subset-specific markers in unstimulated (Ctr-) and SEB-stimulated (Ctr+) CD4^+^T-cells. E Representative stainings showing CD69^+^IFN-γ^+^ cells following stimulation with EBV-but not with MOG/MPB-derived peptides. F. Five untreated PwMS were stimulated with the immunodominant pool of EBV peptides (EBV) and EBV-specific CD4^+^ and CD8^+^T-cells tracked by CD69 and IFN-γ co-expression. Shown are three technical triplicates for each patient (PT). The mean coefficients of variance (CV) and the intraclass correlation coefficients (ICC) are indicated. G. Right: Proliferation of FACS-purified T-cell subsets from healthy donors following stimulation with autologous monocytes and EBV-derived peptides for 7 days according to CellTrace dilution. Shown is one representative Histogram of proliferating Tr1-cells (purified as CD4+CCR5^+^PD1^+^CD27^+^CCR6^−^), bars show mean proliferation of 3 HDs. Right: Cell Trace^−^ CD4^+^T-cells were analysed for GzmK and EOMES expression.

**sFigure 8 Antigen specificities of individual T-cell subsets.**

A. Antigen specificities of T-cell subsets. Reported are frequencies CD69^+^IFN-γ^+^ cells in individual CD4^+^FOXP3^−^T-cell subsets among CD4^+^T-cells and responses in total CD8^+^T-cells in HDs (n=10-21), in untreated (n=10-14) and Natalizumab-treated PwMS (n=17-20). Left panels show responses following stimulation with overlapping peptide pools of MOG and MBP, central panels to an immunodominant peptide pool derived from EBV or and right panels to overlapping peptide pools of VP1 and VP2 from JCV. Squares on top of columns indicate statistical significances as compared to the no peptide control. Stars on top of lines indicate statistical significances between different cohorts. B. Lack of responses of GzmK^+^T-cell subsets, i.e. frequencies of CD69^+^IFN-γ^+^ cells in the indicated subset among total CD4^+^T-cells, to PLP (6 HDs and 6 Natalizumab-treated MS patients) C. CD4^+^CXCR3^+^CCR6^−^“Th1” and CXCR3^−^ CCR6^+”^Th17” cell lines derived from the CSF from 5 different treatment-naïve MS patients were analysed for EOMES expression. Shown are the mean percentages of EOMES^+^ cells, and the p-value is indicated.

**sFigure 9 Specificities of individual T-cell subsets for latent or lytic EBV antigens.**

A. Responses of the indicated T-cell subsets to individual EBV-derived antigens, namely EBNA1, BZLF1 and BMRF1 in HDs (EBNA1: n=9, BZLF1: n=6, BMRF1: n=7), untreated (“MS”, EBNA1: n=9, BZLF1: n=8, BMRF1: n=3 and Natalizumab-treated PwMS (“+NTZ” EBNA1: n=19, BZLF1: n=17, BMRF1: n=11). Responses of FOXP3^+^Tregs (HD: n=7, MS n=3, +NTZ: n=11) were assessed according to CD69 and 4-1BB up-regulation and/or IFN-γ production, all other subsets according to CD69 up-regulation and concomitant IFN-γ production. Squares on top of columns indicate statistical significances as compared to the no peptide control. Stars on top of lines indicate statistical significances between different cohorts. B. Sorted Tr1-like cells, Th1-cells or Tregs from a PwMS were labelled with CellTrace, stimulated with autologous monocytes and EBNA1 peptides and proliferation and CD25 expression analysed after 7 days. C. IL-10 production by Th-cells in response to EBV, EBNA1 and MOG/MBP in Natalizumab-treated PwMS (n=12).

**sFigure 10 Specificities of individual T-cell subsets for EBV antigens in SLE patients stratified according to CNS involvement.**

A. IFN-γ responses of peptide-activated CD4^+^GzmK^+^T-cell subsets, CD4^+^CTL, Th-cells and FOXP3^+^Tregs to EBV and EBNA1 in SLE patients without (CNS-, n=11) or with (n=6) CNS involvement (“CNS +”). B. IL-10 production in EBV and EBNA1-activated EOMES^+^Tr1-like cells and by FOXP3^+^Tregs in SLE patients with and without CNS involvement.

**sFigure 11 Associations of EBNA1-specific T-cells with MS risk factors.**

A. IL-10 production by EOMES^+^Tr1-like cells and FOXP3^+^Tregs, in response to EBV and EBNA1 in PwMS stratified according to HLA-DRB1*15:01 expression. B. Anti-EBNA1 IgG serum levels in HDs (n=15) and MS patients (untreated: n=15, Natalizumab-treated: n=21). C-E: Regression analysis of EBNA1-specific T-cell population with anti-EBNA1 IgG levels. R- and p-values are indicated. C. shows IFN-γ-producing EBNA1-specific cells among total CD4^+^ and CD8^+^T-cells. D. shows EBNA1-specific FOXP3^+^Tregs expressing 41-BB. E. shows EBNA1-specific Th-cells expressing CD40L. F. CD40L up-regulation of CD4^+^T-cell stimulated with EBV- or EBNA1-derived peptides T-cells (n=13). Statistical significance of CD40L expression of Th-cells versus Tr1-cells and Tregs was calculated by two-way Annova.

**sFigure 12: Impact of CD4^+^T-cell subsets on the proliferation of CD8^+^ memory T-cells stimulated with EBV-infected B-cells.**

FACS-purified CD8^+^CD45RA^−^ memory T-cells were labelled with CellTrace and stimulated with EBV-infected B-cells (LCL) and SEB. Shown are the Cell Trace histograms after 7 days in the absence or presence of the indicated CD4^+^T-cell subsets of a representative experiment. The Proliferation indices and the percentage of undivided cells (highlighted in grey) are reported.

**sFigure 13 Negative control for EOMES and IL-10 staining in brain sections.**

Triple immunohistochemistry in bright field of MS brain tissue stained with 5 µg/ml purified rat IgG1 k isotype control (eBRG1), 5 µg/ml purified mouse IgG1k isotype control (P3.6.2.8.1) (both by Thermo Fisher Scientific) and rabbit monoclonal antibody to CD4, performed as described. Only immunoreactivity for CD4+ cells (red) was observed. Bar = 100 µm.

---

| Marker | Fluorochrome | Clone | Brand | cat. N° | Dilution |
| --- | --- | --- | --- | --- | --- |
| **IL-7R** | BB700 | HIL-Ti7R-M21 | BD | 566398 | 30 |
| **CXCR3** | PE-Cy7 | 1C6/CXCR3 | BD | 560831 | 60 |
| **CCR6** | BUV496 | 11A9 | BD | 612948 | 30 |
| **CD4** | BUV563 | SK3 | BD | 612912 | 180 |
| **CD8** | BUV805 | SK1 | BD | 564912 | 200 |
| **L/D dye** | *FVS780* | / | *BD* | *565388* | *200* |
| **CD45RA** | BV605 | HI100 | BD | 562886 | 140 |
| **CD69** | BV786 | FN50 | BD | 563834 | 100 |
| **/** | **/** | **/** | **/** | **/** | **/** |
| **CD3** | BUV737 | UCHT1 | BD | 612750 | 140 |
| **CCR7** | PE-CF594 | 150503 | BD | 562381 | 60 |
| **CD161** | PE-Cy5 | DX12 | BD | 551138 | 40 |
| **CD25** | APC-R700 | 2A3 | BD | 565106 | 40 |
| **CD27** | BV480 | L128 | BD | 566139 | 120 |
| **PD-1** | BV650 | EH12.1 | BD | 564104 | 30 |
| **CCR5** | BUV395 | 2D7/CCR5 | BD | 565224 | 30 |
| **FOXP3** | Alexa Fluor488 | 259D | BioLegend | 320212 | 50 |
| **GzmK** | PE | GM6C3 | SantaCruz | SC56125 | 100 |
| **Eomes** | eFluor 660 | WD1928 | Invitrogen | 50-4877-42 | 25 |
| **GzmB** | BV421 | GB11 | BD | 563389 | 80 |
| **Ki-67** | BV711 | B56 | BD | 563755 | 80 |
| **IFN-γ** | BV750 | B27 | BD | 566357 | 140 |
| **IL-7R** | BB700 | HIL-Ti7R-M21 | BD | 566398 | 30 |
| **CXCR3** | PE-Cy7 | 1C6/CXCR3 | BD | 560831 | 60 |
| **CCR6** | BUV496 | 11A9 | BD | 564659 | 30 |
| **CD4** | BUV563 | SK3 | BD | 566000 | 180 |
| **CD8** | BUV805 | SK1 | BD | 564912 | 200 |
| **L/D dye** | *FVS780* | / | *BD* | *565388* | *200* |
| **CD107a** | BV605 | H4A3 | BioLegend | 328634 | 40 |
| **CD49d** | BV786 | 9F10 | BD | 744588 | 120 |
| **CD29** | BUV661 | 38047 | BD | 750084 | 60 |
| **CD69** | BUV737 | FN50 | BD | 612817 | 40 |
| **T-bet** | PE-CF594 | O4-46 | BD | 562467 | 40 |
| **CD40L** | PE-Cy5 | 24-31 | BioLegend | 310808 | 50 |
| **IL-17A** | APC-R700 | N49-653 | BD | 565163 | 30 |
| **IL-2** | BV510 | 5344.111 | BD | 563265 | 40 |
| **41-BB** | BV650 | H4A3 | BioLegend | 328637 | 40 |
| **TNFα** | BUV395 | MAb11 | BD | 563996 | 160 |
| **FOXP3** | AlexaFluor488 | 259D | BD | 320212 | 50 |
| **GzmK** | PE | GM6C3 | SantaCruz | SC56125 | 100 |
| **Eomes** | eFluor 660 | WD1928 | Invitrogen | 50-4877-42 | 25 |
| **GzmB** | BV421 | GB11 | BD | 563389 | 80 |
| **Ki-67** | BV711 | B56 | BD | 563755 | 80 |
| **IFN-γ** | BV750 | B27 | BD | 566357 | 140 |
| **IL-7R** | BB700 | HIL-Ti7R-M21 | BD | 566398 | 30 |
| **CCR6** | BUV496 | 11A9 | BD | 564659 | 30 |
| **L/D dye** | *FVS780* | / | *BD* | *565388* | *200* |
| **CD8** | BUV805 | SK1 | BD | 564912 | 200 |
| **CD4** | BUV563 | SK3 | BD | 566000 | 180 |
| **FOXP3** | AF488 | 259D | BD | 320212 | 50 |
| **GzmB** | BV421 | GB11 | BD | 563389 | 80 |
| **IFN-γ** | BV750 | B27 | BD | 566357 | 140 |
| **IL-17A** | APC-R700 | N49-653 | BD | 565163 | 30 |
| **Eomes** | PE-Cy7 | WD1928 | Invitrogen | 25-4877-42 | 30 |
| **GzmK** | AlexaFluor647 | G3H69 | BD | 566655 | 40 |
| **CD40L** | BV605 | 24-31 | BioLegend | 310825 | 40 |
| **IL-10** | PE | JES3.9D7 | BioLegend | 501404 | 30 |
| **GM-CSF** | PECF594 | BVD2-21C11 | BioLegend | 562857 | 30 |

