## Supplementary figures and images for "Regulatory T-cells in multiple sclerosis produce IL-10 in the central 1 nervous system but are activated by Epstein-Barr Virus"

### Supplementary Figure 1

Supplemental Figures 1-13

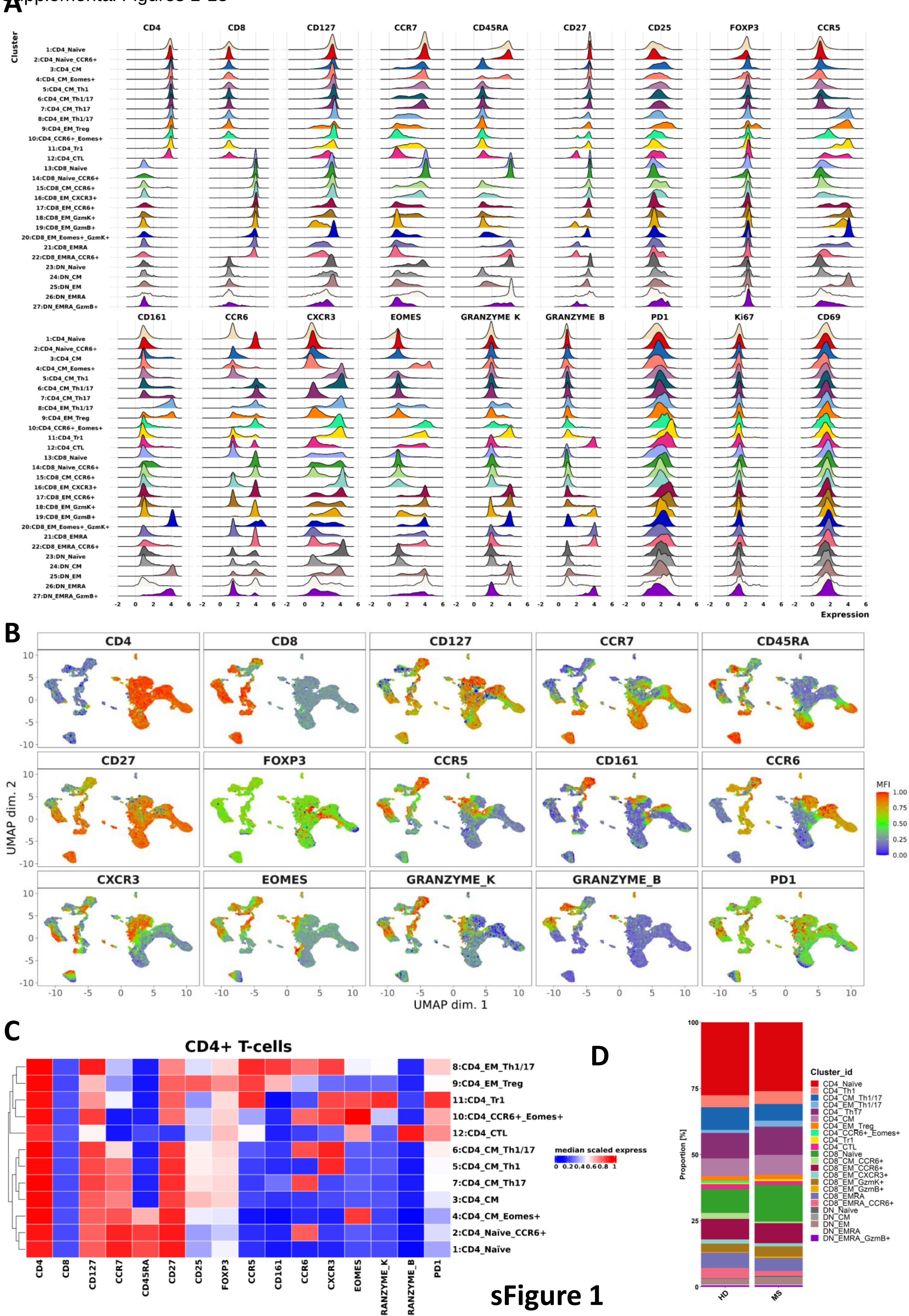


sFigure 1

### Supplementary Figure 2

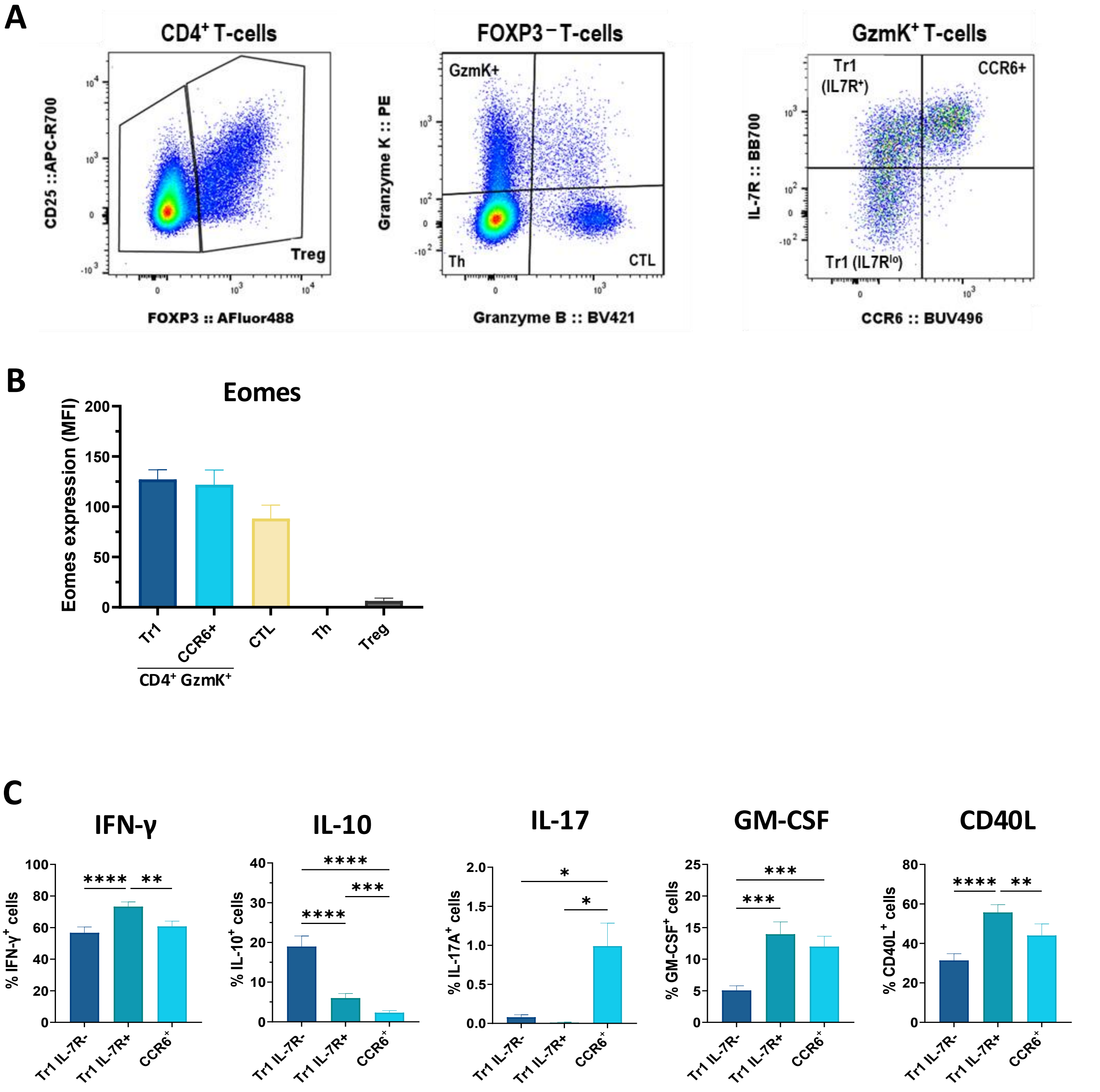

### Supplementary Figure 3

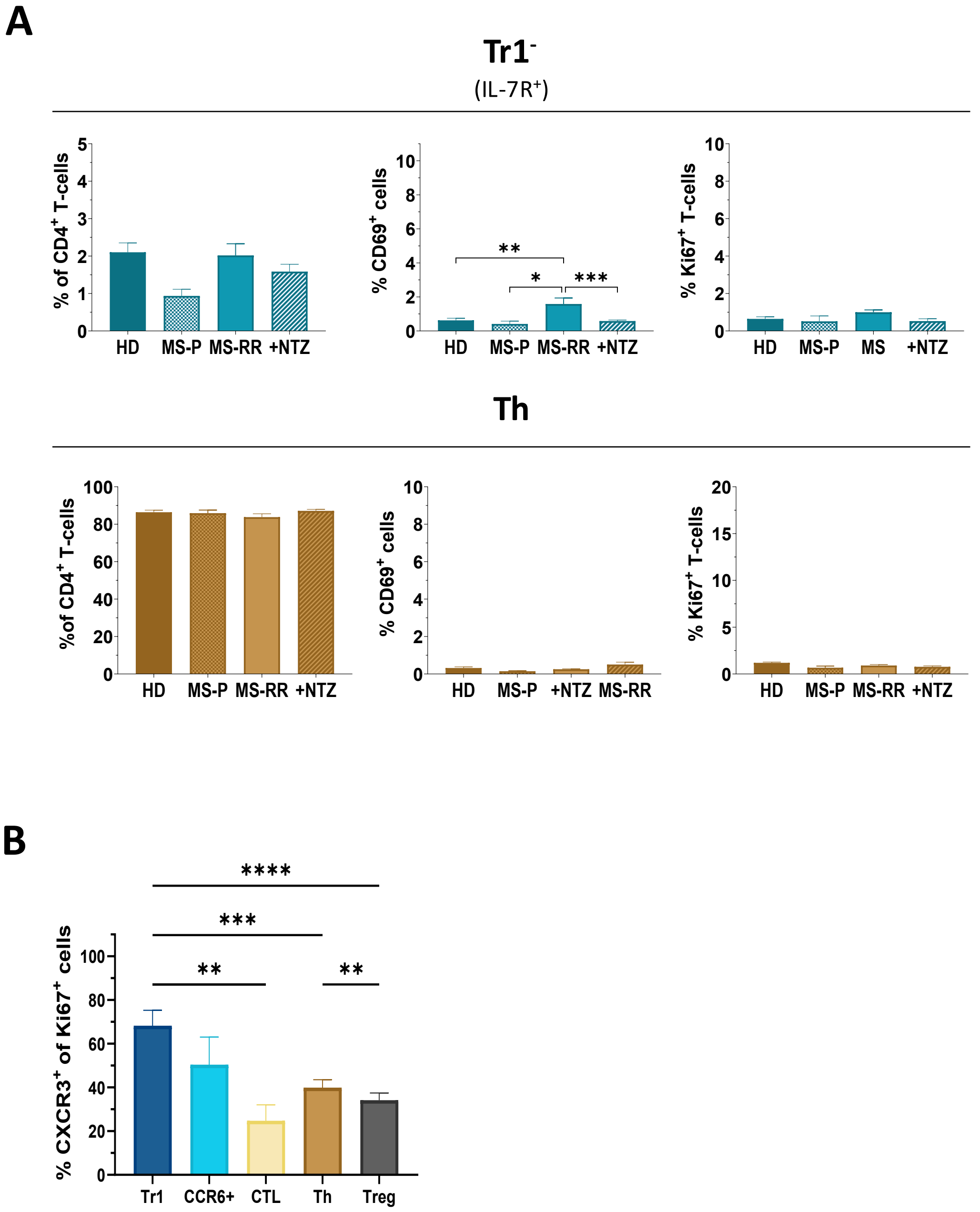

### Supplementary Figure 4

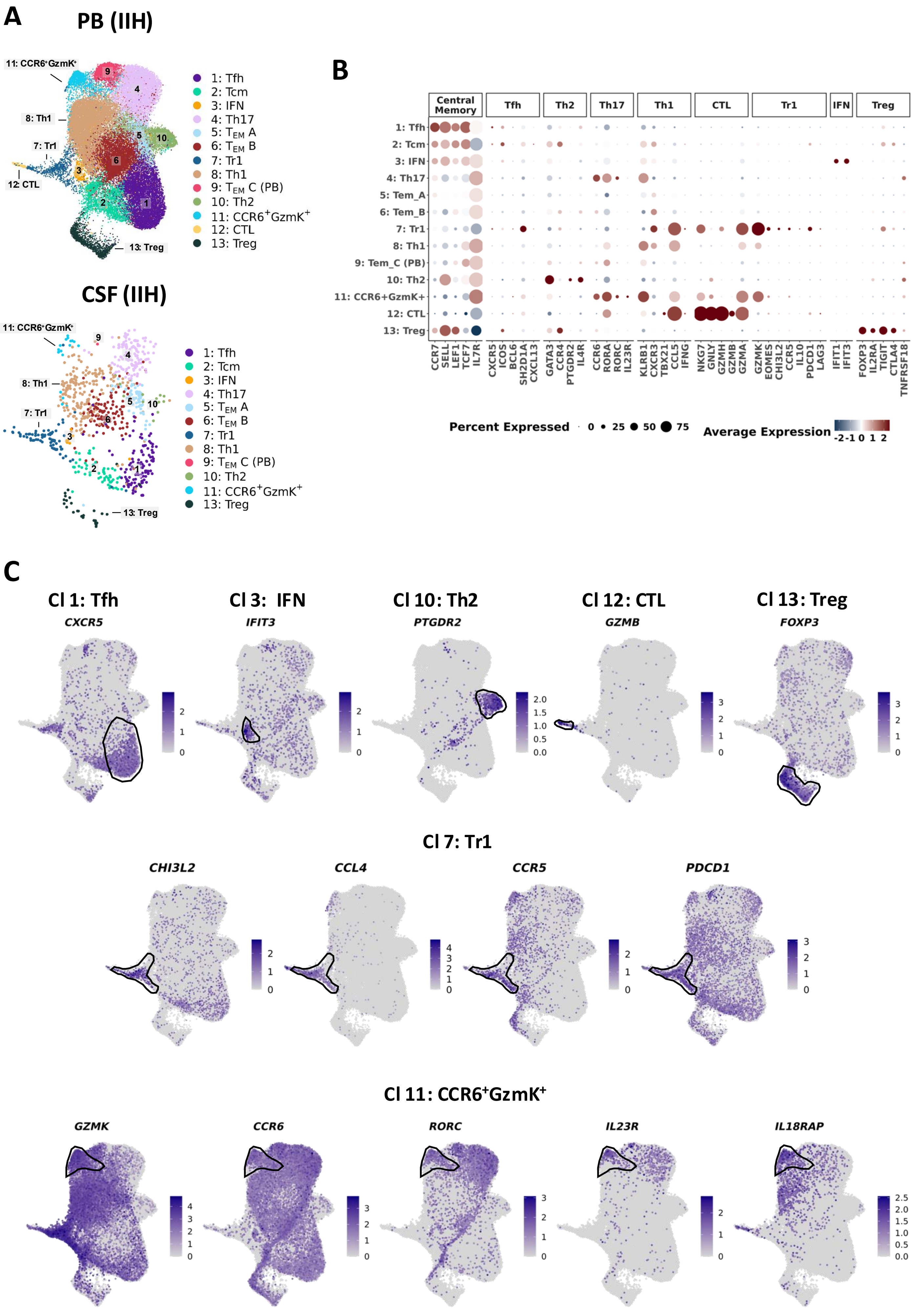

### Supplementary Figure 5

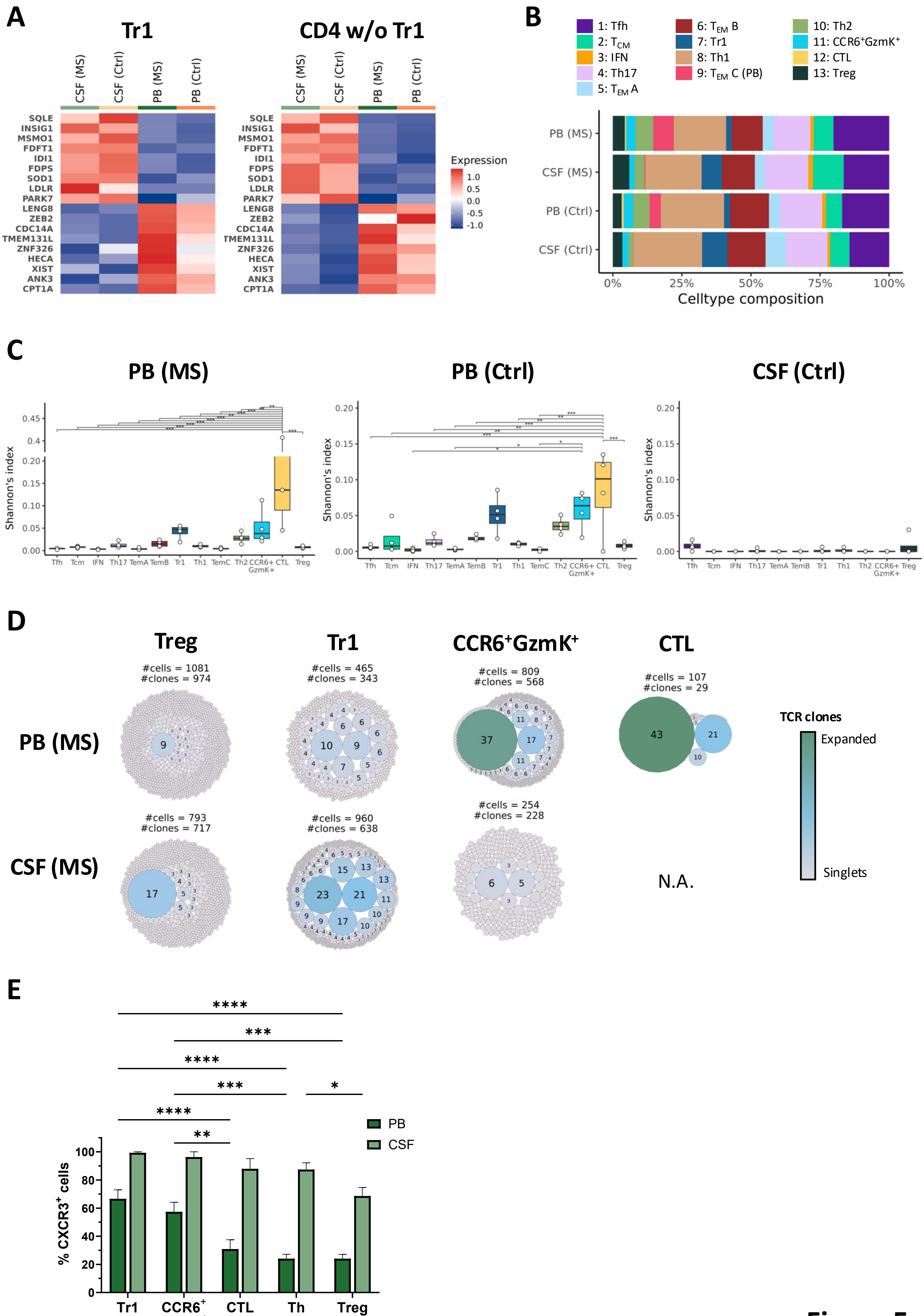

### Supplementary Figure 6

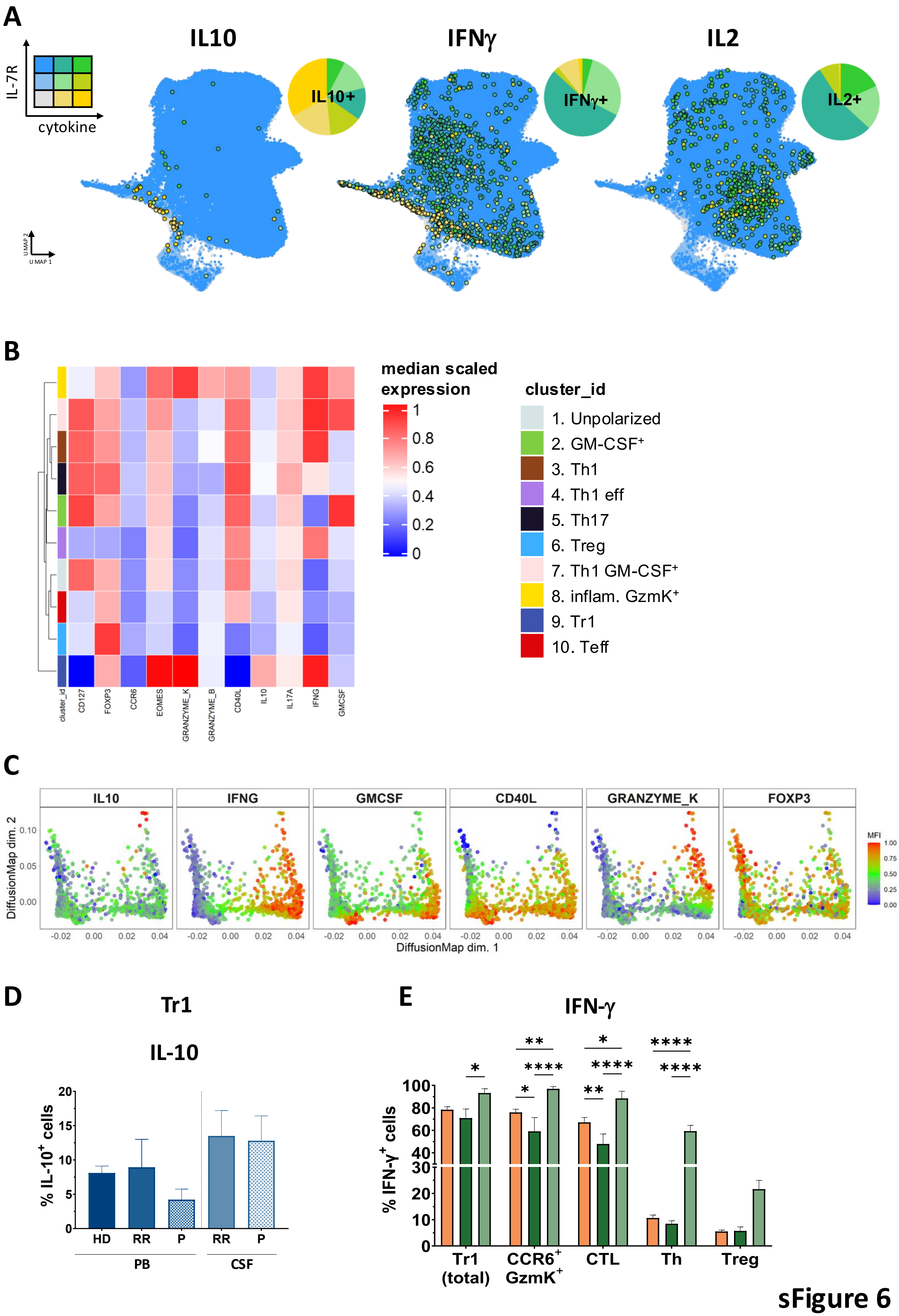

### Supplementary Figure 7

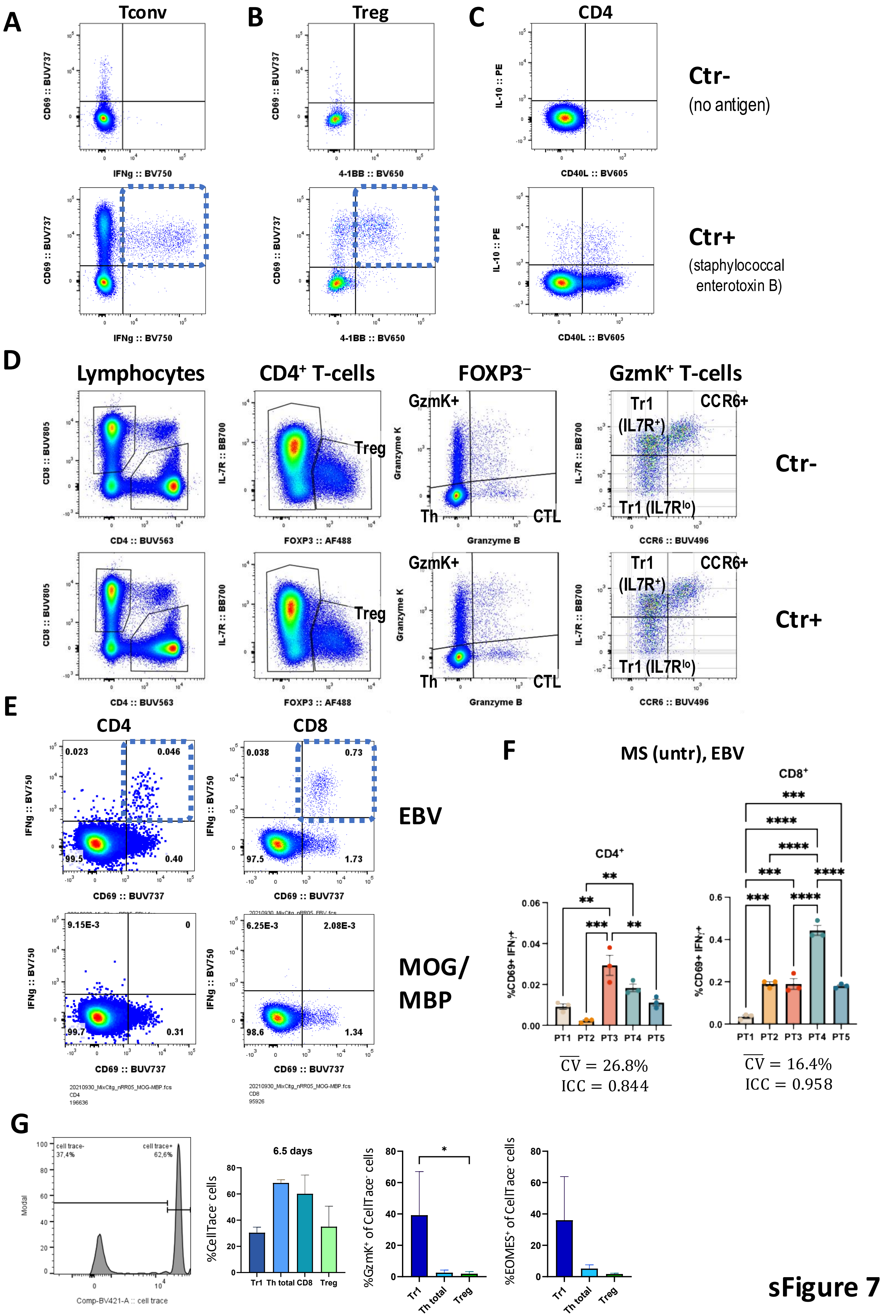

### Supplementary Figure 8

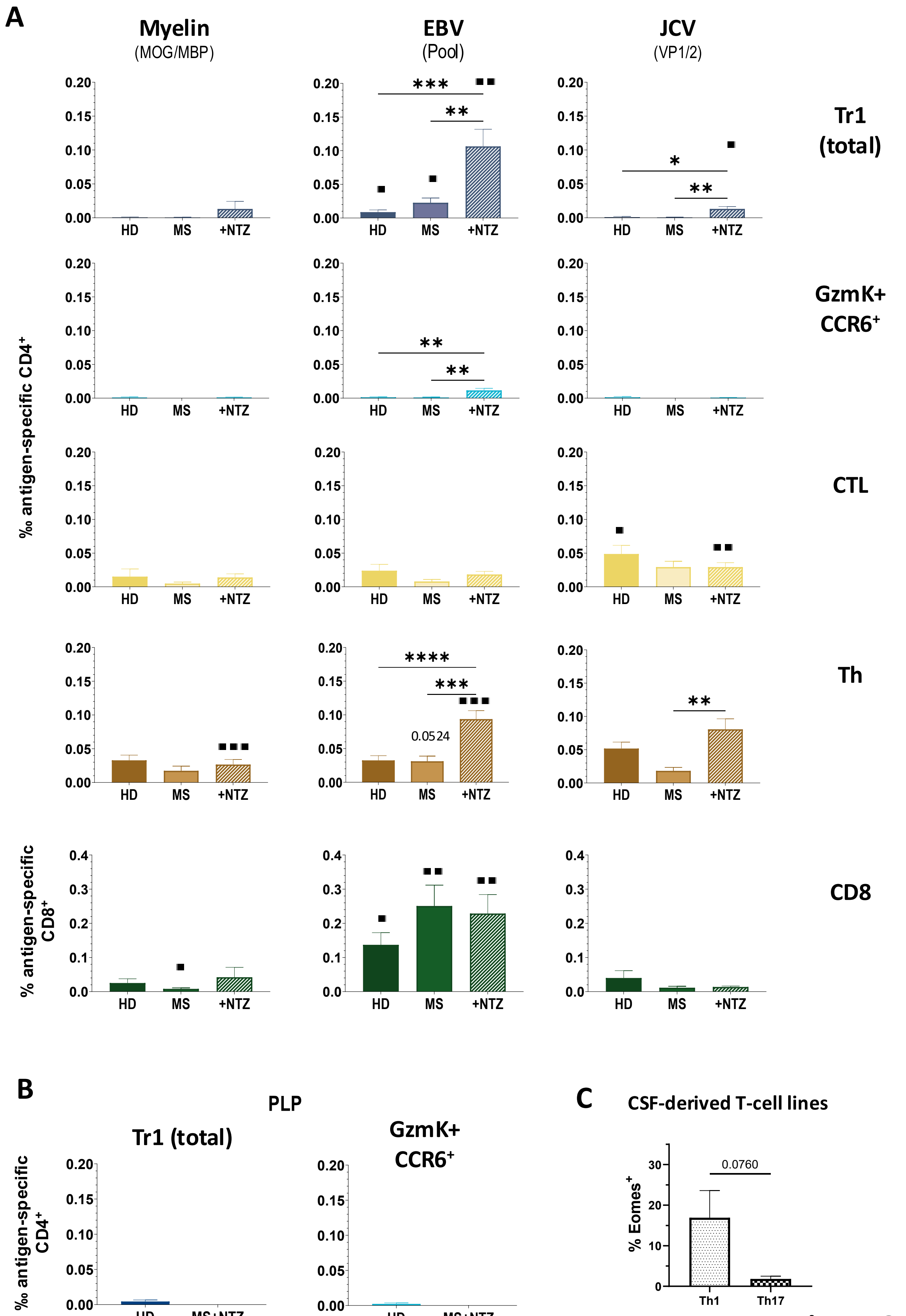

### Supplementary Figure 9

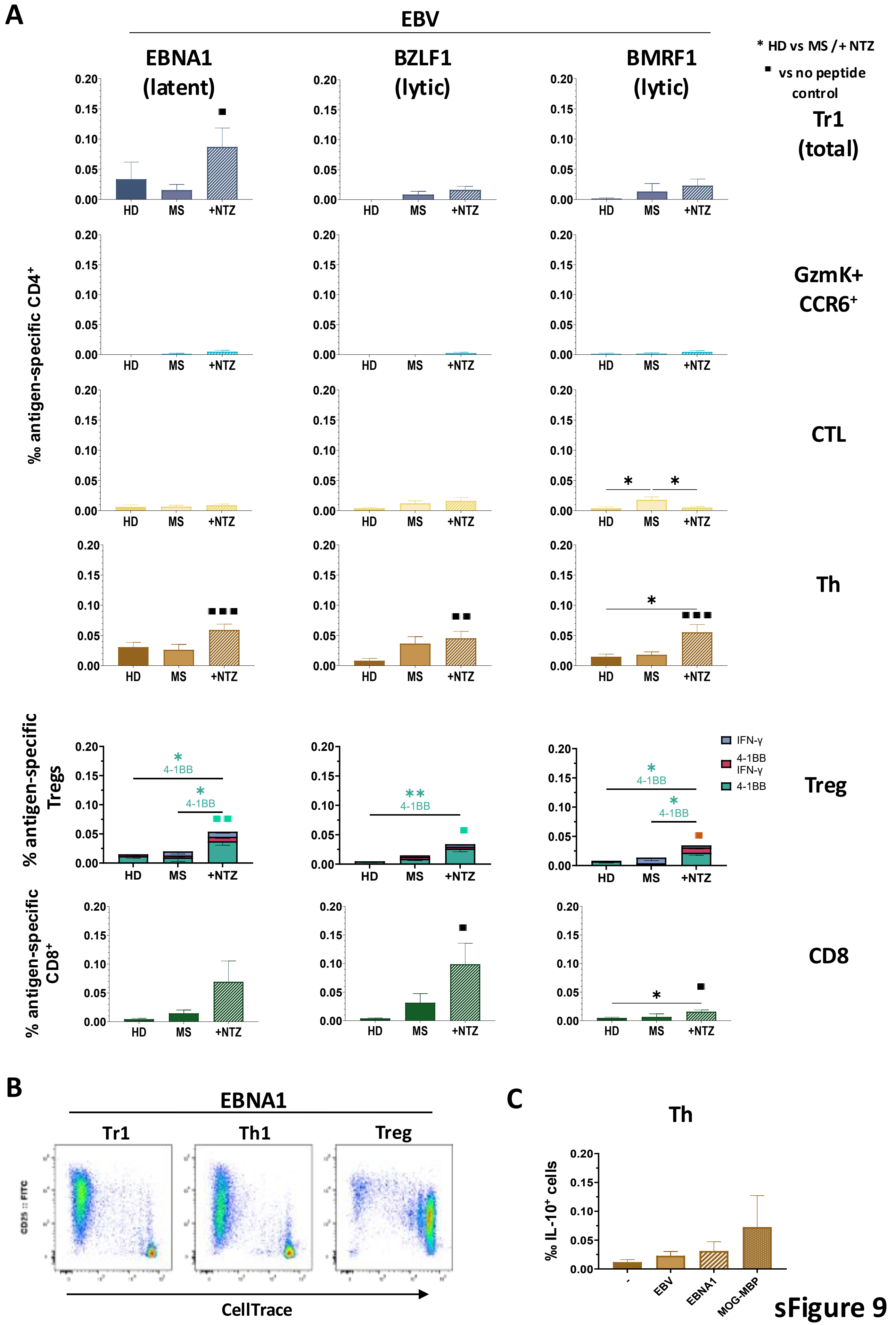

### Supplementary Figure 10

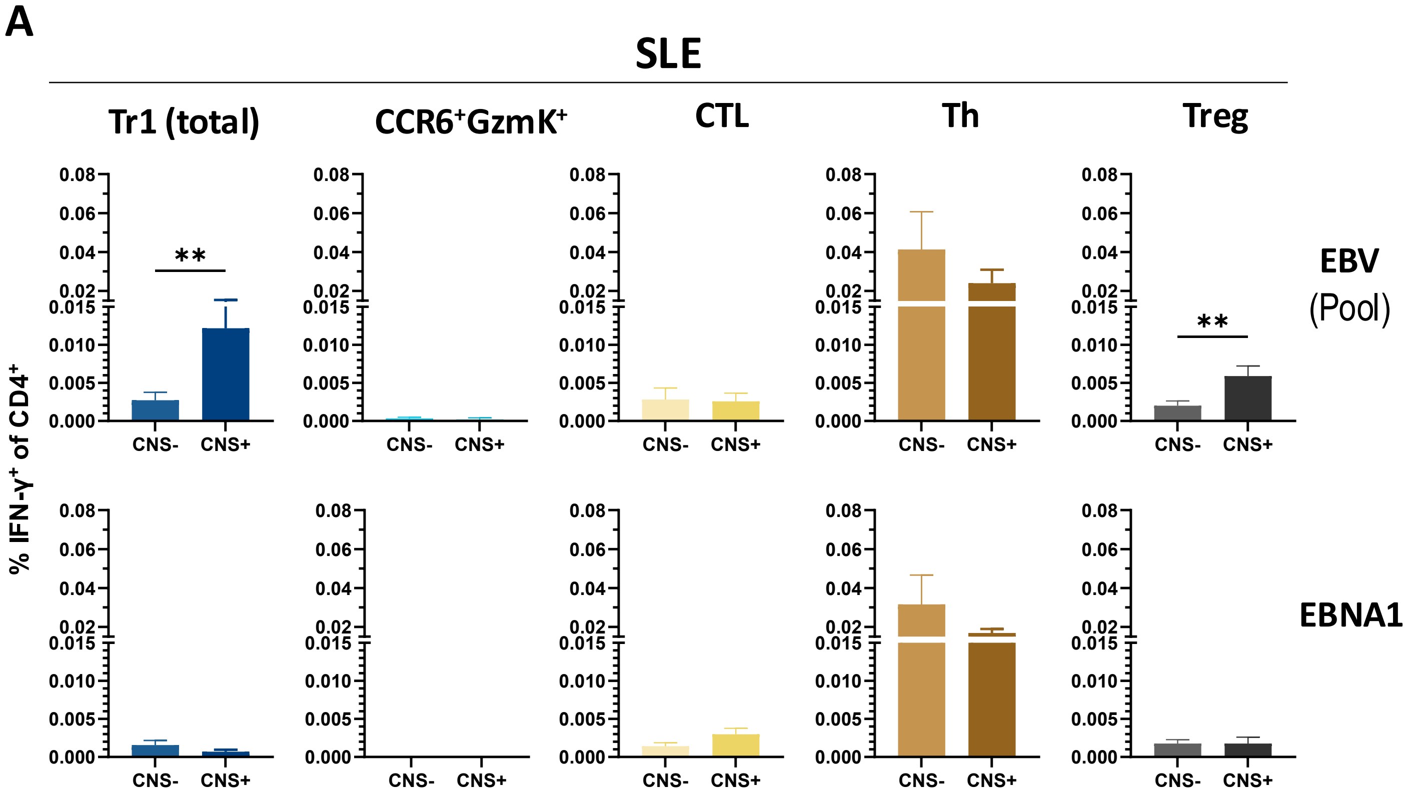

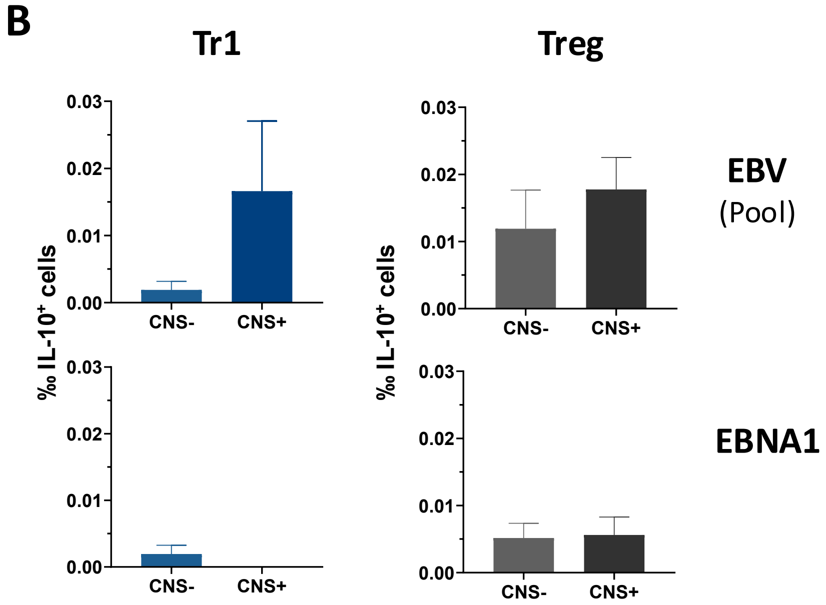

### Supplementary Figure 11

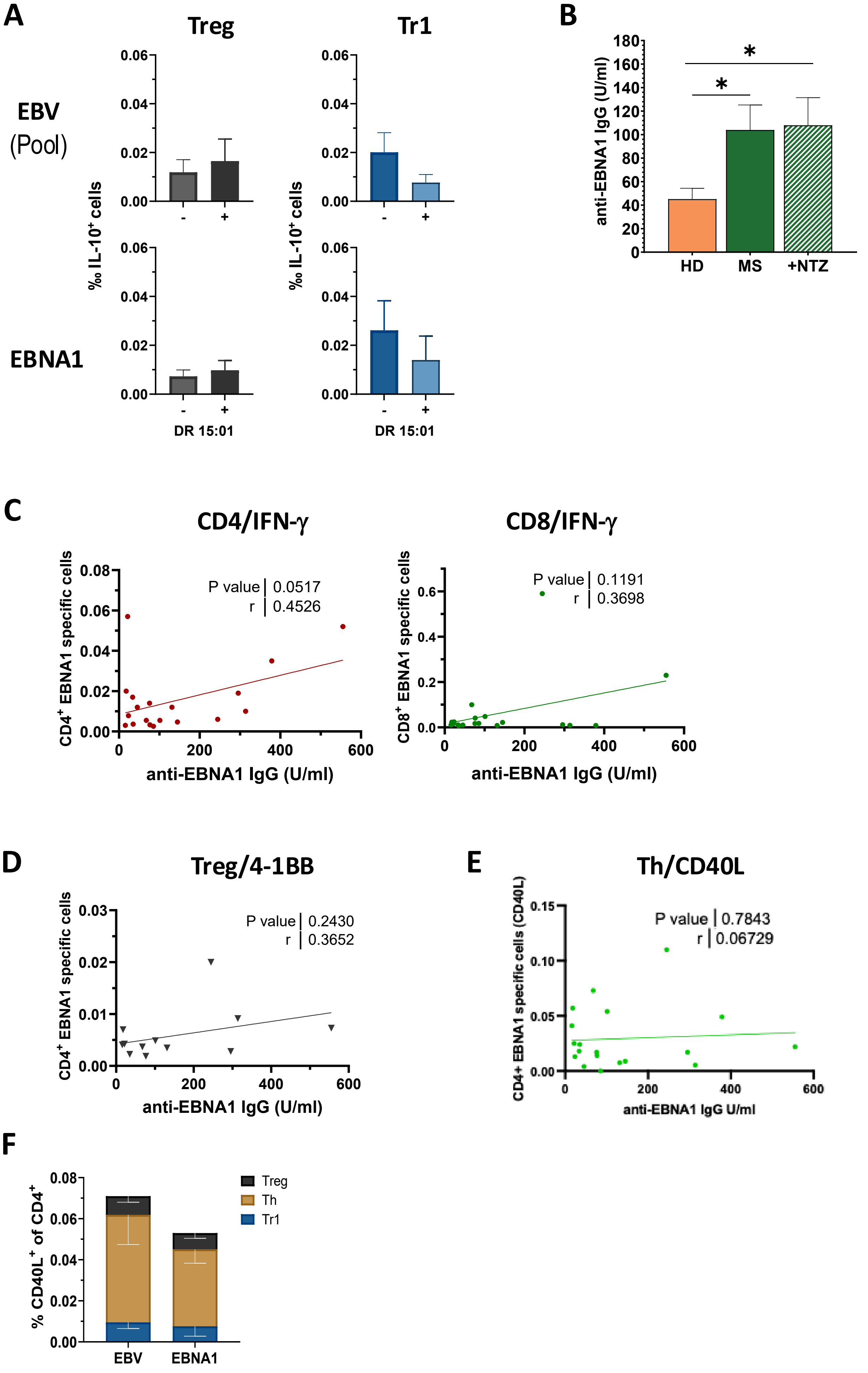

### Supplementary Figure 12

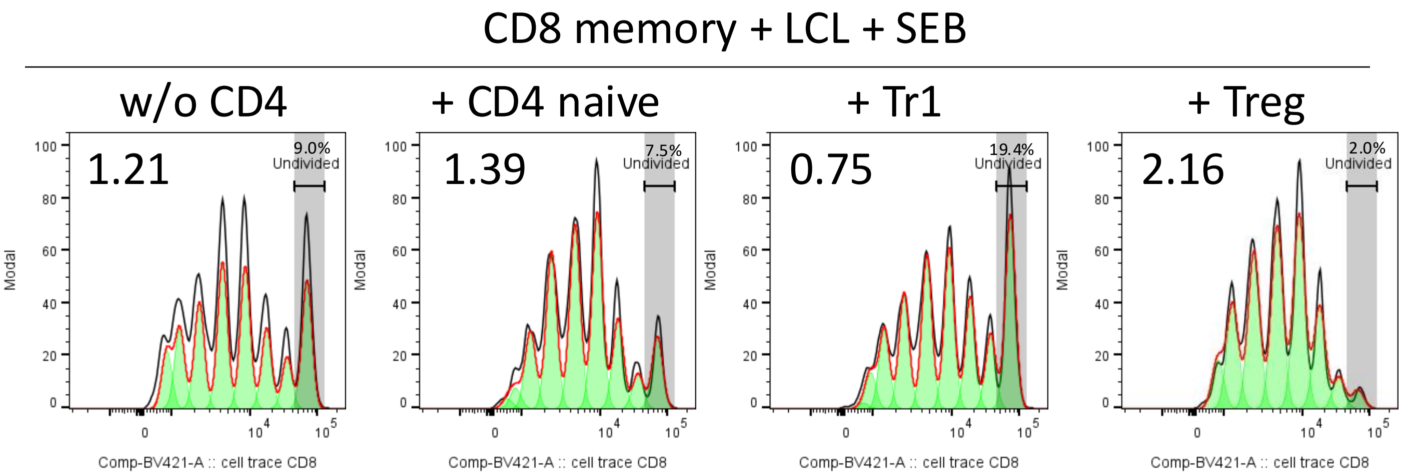

### Supplementary Figure 13

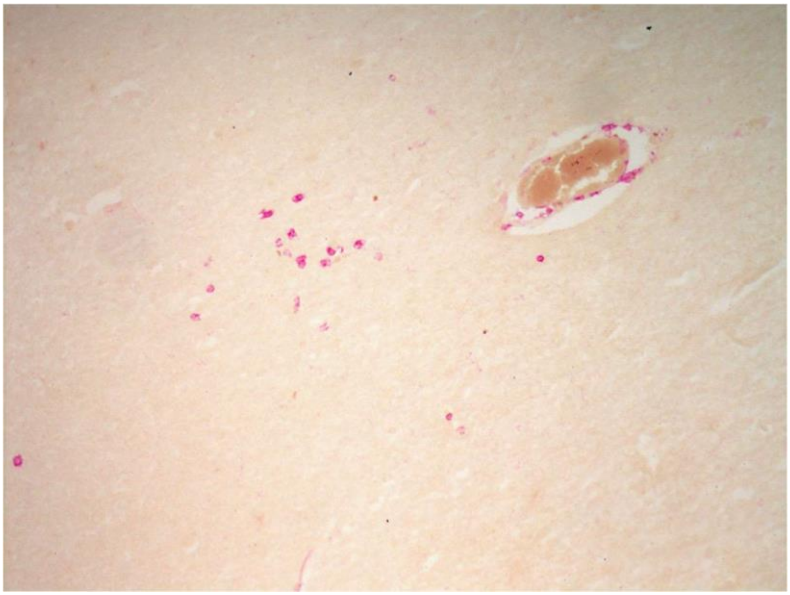

**sFigure 13**
